# Comprehensive Topographical Map of the Serotonergic Fibers in the Mouse Brain

**DOI:** 10.1101/2020.03.18.997775

**Authors:** Janak R. Awasthi, Kota Tamada, Eric T. N. Overton, Toru Takumi

## Abstract

It is well established that serotonergic fibers distribute throughout the brain. Abnormal densities or patterns of serotonergic fibers have been implicated in neuropsychiatric disorders. Although many classical studies have examined the distribution pattern of serotonergic fibers, most of them were either limited to specific brain areas or had limitations in demonstrating the fine axonal morphology. In this study, we utilize transgenic mice expressing GFP under the SERT promoter to map the topography of serotonergic fibers across the rostro-caudal extent of each brain area. We demonstrate previously unreported regional density and fine-grained anatomy of serotonergic fibers. Our findings include: 1) SERT fibers distribute abundantly in the thalamic nuclei close to the midline and dorsolateral areas, in most of the hypothalamic nuclei with few exceptions such as the median eminence and arcuate nuclei, and within the basal amygdaloid complex and lateral septal nuclei, 2) the source fibers of innervation of the hippocampus traverse through the septal nuclei before reaching its destination, 3) unique, filamentous type of straight terminal fibers within the nucleus accumbens, 4) laminar pattern of innervation in the hippocampus, olfactory bulb and cortex with heterogenicity in innervation density among the layers, 5) cortical labelling density gradually decreases rostro-caudally, 6) fibers traverse and distribute mostly within the gray matter, leaving the white fiber bundles uninnervated, and 7) most of the highly labelled nuclei and cortical areas have predominant anatomical connection to limbic structures. In conclusion, we provide novel, regionally specific insights on the distribution map of serotonergic fibers using transgenic mouse.

## 1. INTRODUCTION

Serotonin (5-hydroxytryptamine or 5-HT) is a well-recognized modulator of brain activity and other functions in peripheral organs (Berger et al., 2009). Dysfunction of 5-HT system has been implicated in mood disorders, schizophrenia, addiction, attention deficit hyperactivity disorder, autism spectrum disorders (ASD) and other mental disorders (Lin et al., 2014). In fact, results of several biological studies demonstrate alterations of morphology and/or density of 5-HT neurons in several mental disorders such as depression (Rajkowska et al., 2017), ASD (Makkonen et al., 2008) (Tamada and Takumi, 2015), schizophrenia (Hrovatin et al., 2019), traumatic brain injury (Abe et al., 2016) and in cases like prenatal exposure of psychotropic drugs (Xu et al., 2004) (Maciag et al., 2006) (Weaver et al., 2010), neonatal hypoxia (Reinebrant et al., 2020), or postnatal social isolation (Kuramochi and Nakamura, 2009).

The 5-HT fibers originate from discretely organized cell somas in the raphe nuclei of brainstem and project throughout the brain (Fuxe, 1965) (Vertes and Linley, 2008). In parallel, the activities of these raphe cell bodies are regulated by the monosynaptic inputs from the forebrain and brainstem neurons (Pollak Dorocic et al., 2014). A total of 28,000 5-HT producing neurons in the mouse brain (Ishimura et al., 1988) does not act as a single population, but rather contains parallel sub-systems that differ in connectivity, physiological properties, and behavioral functions (Ren et al., 2018) (Okaty et al., 2019). Its diversity is additionally exhibited by recent transcriptomics study suggesting that the dorsal raphe (DR) 5-HT neurons co-expressing Vglut-3 preferentially innervate the cortex, whereas those co-expressing thyrotropin-releasing hormones innervate the subcortical regions (Ren et al., 2019). The anterograde tracing approaches have revealed that serotonergic efferents from different raphe nuclei to the major brain regions are distinctive and largely non-overlapping (Muzerelle et al., 2016). Although these recent dissections of serotonergic system have unearthed its much finer details, the precise and comprehensive topographical map of total 5-HT efferents in whole brain remains undetermined by the recent neuroanatomical tools.

The serotonergic topography has long been studied with the progressive advent of neuroanatomical techniques such as aldehyde histofluorescence (Fuxe, 1965), autoradiography (Parent et al., 1981), immunohistochemical techniques using antibodies against 5-HT marker enzyme tryptophan hydroxylase (Tph) (Pickel et al., 1977) or 5-HT itself (Steinbusch, 1981). However, they were accompanied by several limitations: These techniques neither allowed the unambiguous identification of labelled 5-HT or non-5-HT neurons, nor were they ideal for visualizing the fine terminal axons due to either the dilution of tracer in highly ramified axons (Parent et al., 1981) or rapid metabolization of released 5-HT, requiring pretreatment before tissue harvesting (Azmitia and Gannon, 1983). Moreover, the synthesis rate or metabolism of 5-HT or Tph, which is subjected to large variability depending on the environmental conditions, may contribute to the alteration in the staining outcome (Nielsen et al., 2006). Therefore, the actual picture of 5-HT innervation in the brain must be more extensive than that revealed by these techniques.

To ensure high fidelity visualization of the fine-grain anatomy of the serotonergic system, we employed SERT (serotonin transporter) -EGFP (enhanced green fluorescence protein) bacterial artificial chromosome (BAC) transgenic mouse in which an EGFP reporter gene is inserted upstream of the coding sequence of the SERT gene (Schmidt et al., 2013). Combined with GFP antibody and high-resolution imaging, we present better organization of serotonergic fibers with the complete visualization of the whole axons, their morphological features and distribution across the whole brain of the adult mice.

## 2. MATERIAL AND METHODS

### 2.1 Animals

All procedures for animal experiments were carried out in accordance with the guidelines of the animal experimentation committee of RIKEN, Japan. The mouse strain used for this research was STOCK Tg (Slc6a4-EGFP) JP55Gsat/Mmucd (RRID: MMRRC_030692-UCD) which was obtained from Research Resources Division, RIKEN. Originally, the mice were obtained from the Mutant Mouse Resource and Research Center (MMRRC) at University of California at Davis, a NIH-funded strain repository, and was donated to the MMRRC by Nathaniel Heintz, The Rockefeller University (Gong et al., 2003). The mice line was maintained by mating male *SERT-EGFP* mice with wild type female C57BL/6J mice (backcrossed more than 5 times). All mice were maintained on a 12/12-hour light/dark cycle and housed in standard plexiglass cages with food and water ad libitum.

### 2.2 Immunohistochemistry

Eleven-week-old male mice (*n*=4) were anesthetized and transcardially perfused with phosphate buffer saline (PBS) followed by 4% paraformaldehyde (PFA) in 0.1 M phosphate buffer (pH 7.4). Brains were removed and immersion-fixed with 4% PFA at 4°C overnight. Coronal (100 μm thick) and sagittal (60 μm thick) sections were made using Leica VT1200 S vibratome (Leica Microsystems, Nussloch, Germany) and collected in PBS containing 0.01% sodium azide and stored at 4°C until use. Immunohistochemical procedures were performed on free floating sections. Sections were treated with 0.3% H_2_O_2_ in PBS for 30 minutes, washed with PBS for 5 minutes three times, incubated with a blocking solution (PBS containing 3% normal goat serum and 0.3% Triton X-100) for 30 minutes at room temperature, and incubated with primary antibodies [Anti GFP rabbit polyclonal antibody (1:500 dilution; #catalog no: A-11122, Thermo Fisher Scientific, Tokyo, Japan)] diluted with the blocking solution overnight at 4°C. Sections were washed with PBS containing 0.3% Triton X-100 (PBST) for 10 minutes four times, incubated with fluorescent-conjugated secondary antibodies [Alexa Fluor 488 Goat anti-Rabbit IgG (1:500 dilution; catalog # A-11034; Thermo Fisher Scientific, Tokyo, Japan)] diluted with the blocking solution for 2 hours at room temperature, washed with PBST for 10 minutes four times, transferred onto Superfrost slides, and mounted with VECTASHIELD with DAPI (Vector Laboratories, Burlingame, CA, USA)

### 2.3 Images acquisition and analysis

The sections were imaged at 10X magnification with VS120 (Olympus, Tokyo, Japan) whole slide scanner fluorescent microscope and FV3000 confocal microscope (Olympus, Tokyo, Japan). The images were processed using ImageJ (Rasband, 1997) and Adobe photoshop CS6, and were converted into grayscale and adjusted for brightness and contrast. Finally, the borders were manually drawn around the nuclei using Inkscape software (Harrington, 2005) upon comparing the SERT immunolabelled sections with its DAPI stained counterparts with the aid of the Allen brain atlas (Allen Institute for Brain Science, 2004). The density of serotonergic fibers in each nucleus was then rated on a scale of 0 to 6 (Table 1).

**Table 1.** Fiber density ratings.

| Density rating | Description | Definition |
| --- | --- | --- |
| 0 | Scanty | lowest density of fibers; more background than fibers |
| 1 | Sparse | low, more background than fibers |
| 2 | Mild | slightly more background than fibers |
| 3 | Moderate | balance between fibers and background |
| 4 | Moderately high | more fibers than background |
| 5 | Dense | possible to distinguish individual fibers |
| 6 | Very dense | highest density of fibers; difficult to distinguish individual fibers |

The representative image for each scale is depicted in supplementary figure 1.

## 3. RESULTS

In this study, we comprehensively mapped the distribution pattern of the serotonergic fibers across the rostro-caudal extent of each brain regions and sub-regions using the SERT-EGFP transgenic mice that was genetically modified to express GFP in the serotonergic fibers. The serotonergic fibers arising from the discrete clusters of cell bodies in the midbrain raphe nuclei innervated nearly all the brain areas in a specific pattern and innervation density. The study plan and the results are depicted schematically in supplementary figure 2. In the following account, we describe the projection pattern and rostro-caudal density gradient in each nucleus of various brain areas and their sub-divisions.

### 3.1 INNERVATION PATTERN IN THE THALAMUS

We analyzed the innervation density and the pattern of SERT-EGFP fibers across the rostro-caudal extent of different thalamic nuclei. No thalamic nuclei were devoid of the innervation. In order to ease the description, we categorized the thalamic nuclei according to the recent classification concept viz; (1) midline and intralaminar nuclei (2) association nuclei (3) principal nuclei (4) epithalamus and (5) the reticular nucleus (Groenewegen and Witter, 2004). Towards the rostral pole of the thalamus, we observed that some of the dorsally diverted collaterals of pathway fibers from the lateral preoptic area (LPO) and lateral hypothalamic area (LHA) passed around the column of fornix and/or along the zona incerta (ZI) to project into the ventrally located thalamic nuclei such as the nucleus reuniens (RE) (Figure 1B & 2C, arrow). Meanwhile, towards the caudal thalamic levels, we noticed that laterally directed collaterals of pathway fibers passed across the ZI to innervate the lateral geniculate (LG) nuclei (Figure 1G). Similarly, rostrally directed collaterals of pathway fibers from the periaqueductal gray (PAG) (Figure 1G), which traversed through the paraventricular thalamic nuclei (PVT) (Figure 1E, arrow), provided the innervation to the dorsally located thalamic nuclei of caudal pole (ex; PVT and other neighboring nuclei).

**Figure 1:**
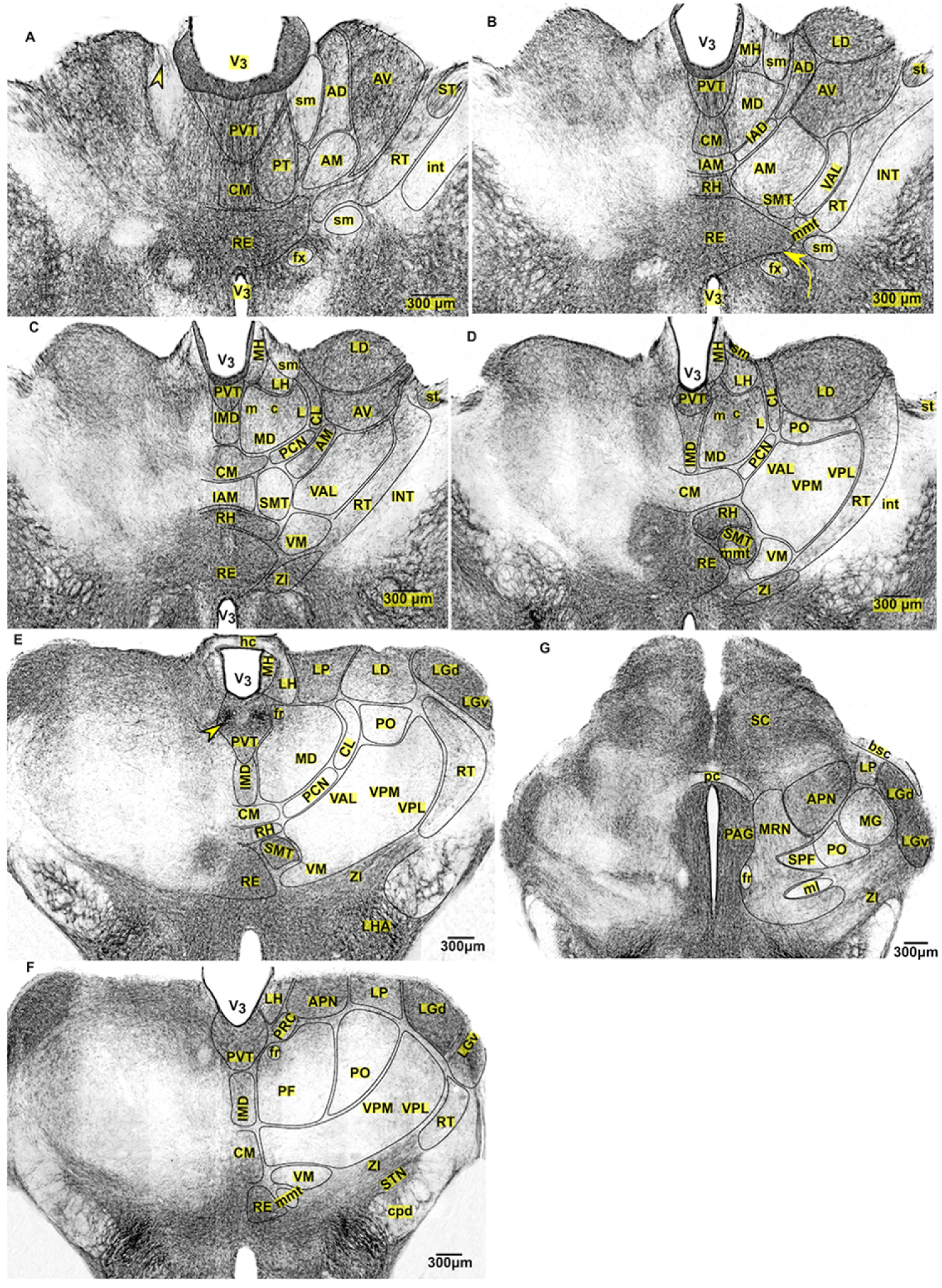
Innervation pattern of SERT-EGFP fibers across the rostro-caudal extent of thalamus. *The terminal labelling density in each nucleus is written numerically within the brackets.* **(A-B)** *Sections from rostral thalamic pole* (**A)** PVT, CM nuclei and nucleus RE (6+). PT and AV nuclei (5+). AD (4+). AM and RT* nuclei (2+). ST nuclei (3+). SM and fx (0), fibers clustered at apex of SM (arrow head). *Rostral most RT was moderately labelled (fig not shown). **(B)** PVT, CM, RH nuclei and nucleus RE (6+). LD and AV nuclei (5+). IAM and AD nuclei (4+). AM and IAD nuclei (2+). VAL and RT nuclei (1+). MD and SMT nuclei (3+). MH (1+). **(C-D-E)** *Sections from mid thalamic level* (**C)** PVT, RH nuclei and nucleus RE (6+). IMD, LD, AV and CL nuclei (5+). CM, PCN and MD nuclei (3+), central part & dorsal area of MD (4+). VAL, VM, RT, SMT nuclei (1+), MH (3+ on ventral area, 1+ on dorsal). **(D)** PVT, RH nuclei, nucleus RE and SMT (6+). IMD, CL and LD nuclei (5+). MH, LH and MD nuclei (3+), central part & dorsal area of MD nuclei (4+). CM, PCN, PO, VAL, VPM, VPL, VM, and RT nuclei (1+) **(E)** PVT*, RE, SMT, LGd & LGv nuclei (6+). LP nuclei (5+). LD nuclei (4+). MH and LH (3+). MD nuclei (2+), less density on its ventral part. IMD and RH nuclei (3+). CM and RT nuclei (1+). PCN, CL, PO, VM, VAL, VPM, and VPL nuclei (<1). *Ascending fiber bundles traversing ahead via PAG were clustered as bilateral bundle in PVT (arrow head). Some of them divert laterally and in-between habenula. **(F-G)** *sections from the caudal thalamic pole* **F)** RE and LGd & LGv nuclei (6+), slightly less fibers on medial part of LGv. PVT nuclei (5+). LH and LP nuclei (4+). PF nuclei (1+; dorsal part, 2+). RT nuclei (1+). PO, VM, VPM and VPL nuclei (<1+). **G)** LGd & LGv nuclei (6+). LP (3+) nuclei. MG nuclei (2+) with slightly higher on lateral part. PO nuclei (1+).

**Figure 2:**
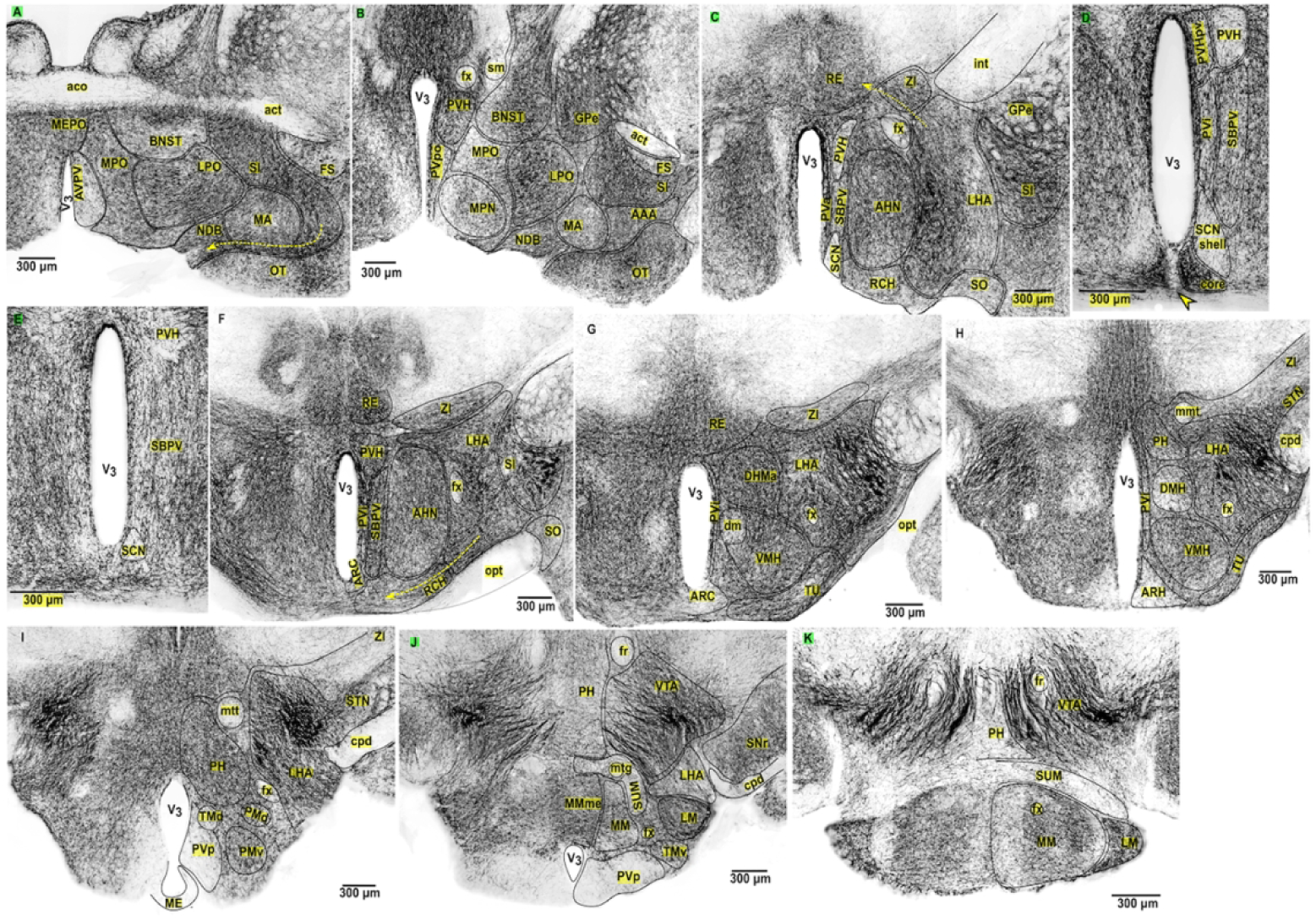
Innervation pattern of SERT-EGFP fibers across the rostro-caudal extent of hypothalamus. **(A-B)** *Sections from preoptic hypothalamus* **(A)** LPO, MPO, MEPO nuclei (5+). AVPV nuclei (2+). Non-hypothalamic nuclei: SI (6+), which contains the collaterals of ascending pathway fibers detached from LPO area. MA nucleus and NDB (5+). Fibers from SI pass medially across the base of MA nucleus and NDB (yellow arrow). OT (5+). Fundus of striatum (FS) contains scattered collaterals of ascending fibers. **(B)** LPO area (6+). PVHap and PVpo nuclei (5+). MPN nuclei (4+). MPO area (4+) with 3+ density on its medial part. Non-hypothalamic nuclei: SI (6+). NDB (5+). MA nucleus (4+). OT (5+) with heavy clustering of fibers over its dorsal area. **(C-F)** *Sections from supraoptic hypothalamus* **(C)** LHA area (6+) with slightly less fibers on lateral part. AHN and ZI (5+). Fibers from LHA pass medially into the thalamus upon traversing through ZI or fx. SBPV, RCH and SO nucleus (3+). PVH; besides the part lining the ventricle (PVHpv; 5+), rest of the PVH is sparsely (1+) labelled. SCN (1+). Non-hypothalamic nuclei: SI (6+). **(D)** SCN; core (6+), shell (1+). Optic fibers entering into SCN were innervated by 5-HT fibers (arrow head). SBPV, PVi nuclei (5+). PVH (1+; except periventricular part, PVHpv 5+). **(E)** SCN (1+). SBPV area (5+). PVH nucleus (1+; except periventricular part, PVHpv 5+). **(F)** LHA (6+). AHN, ZI, RCH, PVH, SBPV, and PVi nuclei (5+). Fibers from LHA pass over supraoptic commissure to enter into the RCH (yellow arrow). Innervation in RCH decreases slightly on medial parts. Dorsally, fibers from LHA enter into the RE of thalamus and into ZI. SO nucleus (3+). Arc nuclei (2+). Non-hypothalamic area: SI (6+) **(G-H)** *Sections from supraoptic hypothalamus* **(G)** LHA (6+). DMHa, TU* nuclei and PVi (5+). VMH nuclei (5+; except dm part, 2+). Arc nucleus (1+). ZI (4+). *Collaterals of pathway fibers were observed to be directly passing into TU nuclei. **(H)** LHA (6+). VMH, TU, PH, STN nuclei, and PVi (5+). ZI (3+). DMH nucleus (2+). Arc nucleus (1+). Some fibers from LHA enter into the rostrally located GP through cpd. **(I-J-K)** *Sections from mamillary hypothalamus* **(I)** LHA (6+). PH and STN nuclei (5+). PMv and PMd (4+). TMd (2+). PVp (1+). ME (0). **(J)** LM* nucleus, and MMme (5+). LHA, MM nuclei, TMv (4+) and PH (3+). SUM (2+). PVp (1+). *Collaterals of pathway fibers from VTA pass laterally to directly enter into the LM nuclei (black segmented arrow). **(K)** LM nucleus (5+). MM nucleus (4+) with mild (2+) innervation on medial part. SUM nucleus (1+)

#### 3.1.1 Midline nuclei

The midline nuclei include paraventricular thalamic (PVT), paratenial (PT), rhomboid (RH) nuclei and nucleus reuniens (RE). We observed that the entire anteroposterior axis of these nuclei was very densely (6+) labeled with a couple of exceptions. For example, the labeling density within RH nuclei decreased to moderate (3+) density at its caudal extent (Figure 1E), while the PT nucleus was densely (5+) labeled (Figure 1A). We found that the ascending pathway fibers traversing ahead from PAG were clustered as two bilateral bundles within the caudal PVT (Figure 1E, arrow) which diffused rostrally to innervate the other adjacent nuclei. There were other nuclei such as the intermediodorsal (IMD), interanteromedial (IAM), and interanterodorsal (IAD), which were located in the midline but were not considered part of the midline nuclei.

#### 3.1.2 Intralaminar nuclei

These nuclei were anatomically associated with the medullary lamina of thalamus, thus were considered as a separate category despite their location in the thalamic midline. It consists of central lateral (CL), paracentral (PC), central medial (CM) and parafascicular (PF) nuclei (Groenewegen and Witter, 2004). We observed that SERT-EGFP fiber density within the CM nuclei gradually decreased from very dense (6+) labelling (Figure 1A) to sparse (1+) density (Figure 1D), and progressively became milder (2+) towards its caudal extent (Figure 1F). We also noticed an intranuclear gradient at its caudal extent (Figure 1F). Similarly, we found that labelling density within CL nuclei gradually decreased from dense (5+) (Figure 1C) to sparse (1+) (Figure 1E) rostro-caudally. Likewise, innervation density within PC nuclei changed from moderate (3+) (Figure 1C) to scanty (0) level (Figure 1E) along the rostro-caudal extent, whereas PF nuclei exhibited intranuclear gradient with milder (2+) density on its dorsal aspect and sparse (1+) ventrally (Figure 1F).

#### 3.1.3 Association nuclei

According to the aforementioned classification, thalamic association nuclei include the anterior nuclei and its subdivisions [anteroventral (AV), anterodorsal (AD), anteromedial (AM), interanteromedial (IAM)], lateral dorsal (LD), lateral posterior (LP), mediodorsal (MD), intermediodorsal (IMD) and submedial thalamic (SMT) nuclei. Among the different subdivisions of the anterior nuclei, we observed that AV nuclei were densely (5+) labelled throughout the rostro-caudal extent (Figure 1A-C). Similarly, AD nuclei received moderately high (4+) projections (Figure 1A-B). AM and IAD nuclei had milder (2+) projections (Figure 1B), whereas the labelling density within IAM nuclei decreased from moderately high (4+) (Figure 1B) to milder (2+) density (Figure 1C) rostro-caudally. We spotted that LD (Figure 1B-E) and LP (Figure 1E-F) nuclei were densely (5+) labeled with slight decrease in their labelling density towards their caudal extent. The density of fibers within the MD nuclei decreased from moderate (3+) to mild (2+) one rostro-caudally (Figure 1 C-E). However, it exhibited higher labelling within the central (c) part compared to its medial (m) and lateral (l) area (Figure 1C-D). Similarly, the innervation density within IMD nuclei decreased from dense (5+) (Figure 1C) to moderate (3+) labelling (Figure 1F) along the rostro-caudal axis. Interestingly, the fiber density within the submedial nucleus exhibited distinct rostral–caudal gradient being moderately (3+) labeled at the rostral half (Figure 1B), sparsely (1+) at the intermediate level (Figure 1C) and its caudal extent was very densely (6+) labelled (Figure 1D-E).

#### 3.1.4 Principal nuclei

Principal nuclei include the ventrobasal (VB) complex [ventral posteromedial (VPM) and ventral posterolateral nuclei (VPL)], ventroanterior lateral complex (VAL), ventromedial nucleus (VM), posterior nucleus (PO), the lateral (LG) and medial geniculate (MG) nuclei (Groenewegen and Witter, 2004). We noticed that the VB, VAL complexes, VM, and PO nuclei received almost no innervation (<1) (Figure 1D-F) except the PO nucleus at its rostral extent, which was sparsely (1+) labelled (Figure 1D). The LG nuclei complex was very densely (6+) innervated throughout the rostro-caudal extent (Figure 1E-G). However, the medial area of the ventral LG nuclei (LGv) had comparably less innervation (Figure 1F). Similarly, we noticed that MG nuclei were mildly (2+) labeled only on its lateral area (Figure 1G), thus showing the intranuclear gradient. We observed that the fascicles of SERT-EGFP fibers arising from the PAG passed laterally along ZI to enrich the innervation of MG and LG nuclei (Figure 1G).

#### 3.1.5 Medial and lateral habenula (epithalamus)

We found that the habenula was less innervated compared to other dorsally located thalamic nuclei. Medial habenula were sparsely (1+) labelled towards the rostral pole (Figure 1B). As traced caudally, the whole area of lateral habenula and only the lateral part of the medial habenula were moderately (3+) labelled (Figure 1C-D). This pattern was due to the collaterals of pathway fibers ascending ahead through the PVT being directed laterally along the borders in-between the medial and lateral habenula. The labelling density within the lateral habenula was moderately high (4+) towards the caudal extent (Figure 1F). Overall, the lateral habenula was innervated in higher density compared to the medial habenula.

#### 3.1.6 Reticular nucleus

Except the moderately (3+) labelled rostral pole of reticular nuclei (Figure 1A), the rest of its extension received sparse (1+) innervation (Figure B-F). However, the innervation density was slightly higher than adjacently located ventrobasal complex of the thalamus (Figure 1D).

#### 3.1.7 Other areas

The white fiber bundles; viz. fasciculus retroflexus (fr), mamillotegmental tract (mmt), posterior commissure (pc), internal capsule (int) and cerebral peduncle (cpd) were devoid of the innervation. However, few SERT-EGFP fibers passed around the white fibers within the cerebral peduncle (Figure 1F) to reach the globus pallidus (GP). We observed the collaterals of pathway fibers travelling within the stria medullaris (sm) and stria terminalis (st). Within stria terminalis, the fibers were dispersed throughout the tract (Figure 6D). However, fibers were clustered only in the dorsal part of sm leaving the rest of the part being devoid of the innervation (Figure 1A, arrow head).

### 3.2 INNERVATION PATTERN IN THE HYPOTHALAMUS

We categorized hypothalamic nuclei according to a conventional classification method based on regional location, viz; preoptic region, supraoptic region, tuberal region and mammillary region (D. G. Stuart, 1962). Preoptic and supraoptic regions were located towards the rostral hypothalamic zone, and the tuberal region was situated in the middle hypothalamic zone, whereas the mamillary region was positioned in the caudal hypothalamic zone. We noticed that the hypothalamic nuclei were heterogeneously innervated.

#### 3.2.1 Preoptic region (Figure 2A-B)

The ascending forebrain bundle traversed rostrally through lateral preoptic region (LPO). Few thin, punctate fibers of innervation were scattered in between the thick ascending pathway fibers. Some of the detached fibers from the main ascending bundle were clustered separately within the substantia innominata (SI). The median preoptic nucleus (MEPO) and rostral part of medial preoptic area (MPO) were densely (5+) innervated (Figure 2A). However, the labelling density in MPO decreased as traced caudally, exhibiting the intranuclear gradient. The fibers were distributed moderately high (4+) on its lateral area and moderately (3+) on its medial part. (Figure 2B). The anteroventral periventricular nuclei (AVPV) were mildly (2+) innervated (Figure 2A), whereas the medial preoptic nucleus (MPN) received moderately high (4+) projection (Figure 2B). The other non-principal hypothalamic nuclei present in the preoptic region; viz, magnocellular nuclei (MA) and nucleus of diagonal band (NDB) were densely (5+) innervated (Figure 2A), although the labelling density in MA decreased slightly as traced caudally (Figure 2B). We observed that the collaterals of pathway fibers which were clustered in SI diverted medially to run over the lower border of MA and NDB in the area adjoining to OT, to finally enter into the septal nuclei (Figure 2A, arrow).

#### 3.2.2 Supraoptic region (Figure 2C-F)

The ascending forebrain bundle traversed rostrally through the lateral hypothalamic area (LHA) with few interspersed fibers of innervation. The collaterals of ascending bundle passed dorsally into the thalamic RE (Figure 2C, arrow) and laterally into the ZI (Figure 2F). Some of the collaterals ran ventromedially through supraoptic commissure and retrochiasmatic nuclei (RCN) (Figure 2F, arrow) to project into the rostrally located suprachiasmatic nuclei (SCN) (Figure 2D). We found that the supraoptic nuclei (SO) (Figure 2C & F) and rostral part of RCN (Figure 2C) were moderately (3+) labelled. However, the labelling density of latter increased densely (5+) caudally (Figure 2F). Anterior hypothalamic nuclei (AHN) were densely (5+) innervated throughout the rostro-caudal extent. Interestingly, we noticed that SCN exhibited both intranuclear and rostro-caudal gradients. Rostral (Figure 2C) and caudal (Figure 2E) SCN was sparsely (1+) innervated. However, the intranuclear gradient was observed in midway SCN (Figure 2D). Its ventromedial area (core) received very dense (6+) projections making it one of the most heavily innervated brain areas while the dorsolateral area (shell) was sparsely (1+) innervated. Similarly, we observed rostro-caudal gradient in the sub-paraventricular zone (SBPV) in which innervation density increased from moderate (3+) (Figure 2C) to dense (5+) levels (Figure 2F). Paraventricular hypothalamic nuclei (PVH) exhibited both intranuclear and rostro-caudal gradients. Rostrally, except its densely (5+) innervated periventricular part (PVHpv) (Figure 2D), the rest of its area was sparsely (1+) labelled (Figure 2C-E). However, at its caudal extent the whole PVH nucleus was densely (5+) labelled (Figure 2F).

#### 3.2.3 Tuberal region (Figure 2G-H)

The ascending forebrain bundle traversed rostrally through the perifornical LHA of tuberal region. The ventromedial hypothalamic nuclei (VMH) exhibited intranuclear gradient across the rostro-caudal axis. Rostrally, its dorsomedial (dm) part was mildly (2+) innervated compared to the rest of the nucleus (5+) (Figure 2G). However, the fibers were densely (5+) and homogeneously distributed throughout the nucleus as traced caudally (Figure 2H). Similarly, we noticed that the dorsomedial hypothalamic nuclei (DMH) were densely (5+) innervated rostrally (Figure 2G) and labelled mildly (2+) at its caudal extent (Figure 2H), displaying the rostro-caudal gradient. The arcuate nucleus exhibited sparse labelling (1+) throughout the antero-posterior axis. We found that the collateral fibers arising from the main ascending bundle in LHA projected ventrally into the tuberal (TU) nuclei making it densely (5+) innervated.

#### 3.2.4 Mamillary region (Figure 2I-K)

We observed that the innervation density within the posterior hypothalamic nuclei (PH) decreased from dense (5+) (Figure 2H-I) to sparse (1+) density (Figure 2K) rostro-caudally. Moreover, we noticed that PH nuclei served as the route for the collateral fibers (arising from the main ascending bundle in the LHA) to pass dorsally into the thalamus (Figure 2H). Similarly, the posterior part of periventricular nuclei (PVp) (Figure 2I-J) and supramamillary nuclei (SuM) (Figure 2J, 2K) had sparse (1+) labelling throughout the rostro-caudal extent. Likewise, the both ventral and dorsal parts of premamillary (PMv, PMd) nuclei were labelled in moderately high (4+) density (Figure 2I). We spotted collaterals arising from the ascending fiber bundle at VTA running directly into the lateral mamillary nuclei (LM) making it densely (5+) innervated (Figure 2J, arrow). Similarly, the ventral part of tuberomammillary (TMv) was labelled in moderately high (4+) density (Figure 2J), whereas the dorsal part being mildly (2+) innervated (Figure 2I). The median part of medial mamillary nuclei (MMme) was densely (5+) labelled, while the rest had innervation with slightly less density (4+) (Figure 2J). However, the pattern was in contrary to its caudal extent (Figure 2K). Strikingly, the median eminence had almost no SERT-EGFP labelled fibers (Figure 2H-I).

There were few nuclei which extended over more than one region and the patterns they exhibited were as follows: the mediolaterally extended nuclei ZI carried the collaterals of the main ascending bundle from LHA towards laterally located thalamic reticular nuclei or the geniculate nuclei (Figure 1E-G). We observed that the SERT-EGFP fiber distributed over ZI gradually decreased from the dense (5+) (Figure 2F) to sparse (1+) density (Figure 2I) rostro-caudally. Subthalamic nuclei (STN) that were located just above the cerebral peduncle (cpd) were densely (5+) labelled because of the directly entering collaterals of ascending bundle fibers from LHA (Figure 2H-I). We noticed heterogenicity among the different subdivisions of periventricular nuclei (PV). The preoptic (PVpo) (Figure 2B), anterior (PVa) (Figure 2C), and intermediate (PVi) (Figure 2D) parts of PV nuclei were densely (5+) innervated. The AVPV and PVp nuclei with less innervation have been already included in the preoptic region and mamillary region, respectively.

#### 3.2.5 Other areas within the hypothalamic zone

The optic tract (opt) was completely devoid of innervation (Figure 2F-G). However, some fibers which entered into the SCN received serotonergic innervation (Figure 2D, arrow head). The white fiber bundles, viz. fornix (fx) (Figure 2B), mamillotegmental tract (mtt) (Figure 2H-I), anterior commissure (aco, act) (Figure 2A, B) and cerebral peduncle (cpd) (Figure 2 I-J) appeared almost empty with very few scattered fibers traversing along or through it.

### 3.3 INNERVATION PATTERN IN THE AMYGDALA

We observed that no nuclei within the amygdaloid complex were devoid of the serotonergic innervation. However, they exhibited heterogenicity in the innervation density and gradient along the rostro-caudal axis. We noticed that collaterals of pathway fibers entered into the amygdala via two routes, via the lateral hypothalamic area (LHA) / substantia innominata (SI) (Figure 3A, arrow) and via the stria terminalis (st) (Figure 3C-D). The fibers contained in stria terminalis originated from the main ascending bundle at lateral preoptic area (LPO) (Figure 6D) and ran caudally over the thalamus to ultimately project to the amygdalar nuclei (Figure 3C, D). These collaterals of pathway fibers were scattered among the thin and punctate fibers of innervation and were difficult to trace further. We classified the amygdaloid nuclei into three groups: (1) the deep or basolateral group (2) the superficial or cortical-like group and (3) the centromedial group (McDonald, 1998). There were other nuclei which did not easily incorporate into any of these groups, which we describe separately.

**Figure 3:**
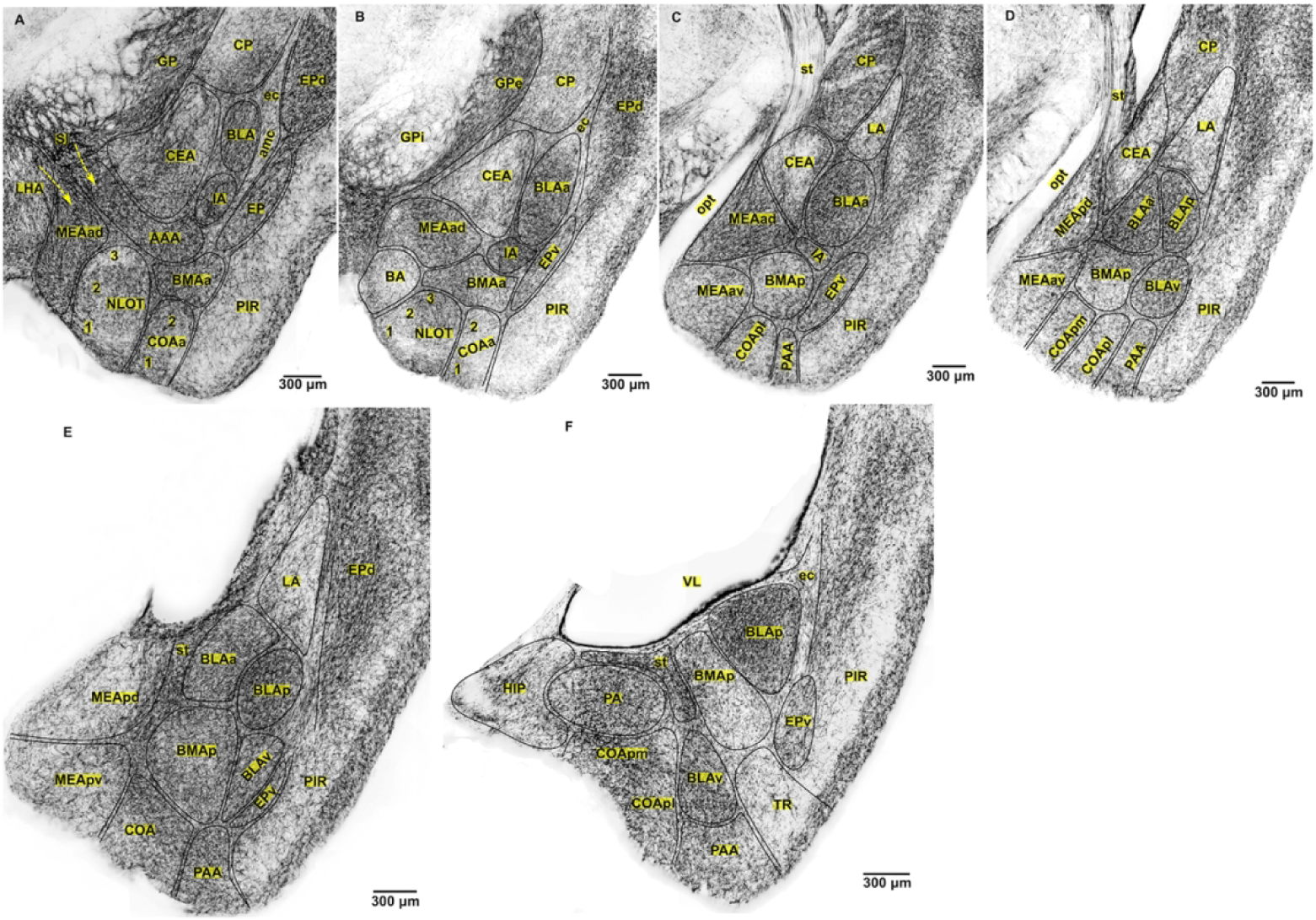
Innervation pattern of SERT-EGFP fibers across the rostro-caudal extent of amygdala. **(A-B)** *Rostral amygdalar section* **(A)** MEAad, medial part of AAA (6+). Lateral part of AAA, IA, BLA and BMAa nuclei (5+). CEA, NLOT*, and COAa (4+). *Rostral NLOT (5+) (figure not shown). Non amygdalar nuclei: EPd (6+) and EPv (5+). Fibers enter into the amygdala from LHA and SI (yellow arrow). **(B) \***BLAa, IA (6+). MEAad, and BMAa nuclei (5+). NLOT-layer 1&3 (4+), layer 2 (1+). CEA and BA nuclei (3+). COAa (2+). Non amygdalar nuclei: EPd (6+) EPv (5+). * BLAa was bounded by very dense clustered fibers within the external capsule on medial and ventral side. **(C-D)** *Mid amygdalar section* **(C)** BLAa (6+). MEAad, and IA nuclei (5+). LA, CEA, MEAav, BMAp, COApl, and PAA (3+). Fibers travelling within stria terminalis (st) project into the CEA and MEAad nuclei. **(D)** BLAa and BLAp (6+). BLAv nucleus (5+). BMAp nucleus (4+), shows the intranuclear gradient. LA, CEA, MEApd, MEAav, COApm, COApl, and PAA (3+). Fibers travelling within stria terminalis (st) project into the CEA and MEApd nuclei. **(E-F)** *Caudal amygdalar section* **(E)** BLAp nuclei (6+). BLAa, BLAv, BMAp, COA, and PAA (5+). MEApd, MEApv, and LA nuclei (3+). Stria terminalis (st) was very densely (6+) clustered with SERT-EGFP fibers both in (E) and (F). **(F)** BLAp nuclei (6+). PA, BLAv, COA (pm,pl), and PAA (5+). BMAp nucleus (3+). Transition zone (TR) between PAA and PIR was mildly (2+) innervated.

#### 3.3.1 Deep or basolateral group

According to the aforementioned classification, the basolateral group includes the lateral amygdalar nucleus (LA), basolateral amygdalar nucleus (BLA), and basomedial amygdalar nucleus (BMA). LA nuclei were moderately (3+) labelled throughout the rostro-caudal extent, though the density in its apical portion was less compared to the basal area (Figure 3C-E). We noticed that BLA nuclei were among the most heavily innervated nuclei (Figure 3A-F). Among its different subdivisions, both the anterior (BLAa) and posterior (BLAp) nuclei were very densely (6+) labelled. However, the fiber density within BLAa decreased slightly at its caudal extent. The ventral part (BLAv) was densely (5+) innervated (Figure 3D-E). Similarly, we observed variations in the innervation density among the different subdivisions of BMA nuclei. Its anterior part (BMAa) was densely (5+) labelled (figure 3A-B). However, the innervation density in the posterior part (BMAp) increased from moderate (3+) (Figure 3C) to dense (5+) (Figure 3E) level and then decreased again towards its caudal extent (Figure 3F).

#### 3.3.2 Superficial or cortical-like group

Superficial or cortical-like group of amygdalar nuclei is also known as corticomedial nuclei. It consists of nucleus of lateral olfactory tract (NLOT), bed nucleus of accessory olfactory tract (BA), anterior and posterior cortical amygdalar nucleus (CoAa and CoAp), and the piriform-amygdaloid area (PAA). We noticed that NLOT exhibited change in both the innervation pattern and density along the rostro-caudal axis. It was densely (5+) innervated at its rostral pole without a laminar pattern (fig not shown). At its midway along the rostro-caudal axis, it appeared distinct because of slightly less innervation (4+) compared to its surrounding area (Figure 3A). We observed a tri-laminar pattern at its caudal extent with a less innervated intervening layer (layer 2) compared to the rest of the layers (4+) (Figure 3B). Similarly, BA was moderately (3+) labelled (Figure 3B). We also observed that the innervation density within CoAa decreased from moderately high density (4+) to milder (2+) level (Figure 3A-B), whereas the density within CoAp increased from moderate (3+) (Figure 3C) to dense (5+) (Figure 3F) labelling pattern along the rostro-caudal axis. Similarly, fiber density within the PAA also increased from moderate (3+) to dense (5+) pattern (Figure 3C-F) rostro-caudally.

#### 3.3.3 Centromedial group

The centromedial group consists of medial (MeA) and central amygdalar (CeA) nuclei. CeA nuclei were moderately (3+) labelled throughout the anteroposterior extent (Figure 3B-D) except at its moderately high (4+) labelled rostral most end (Figure 3A). We observed that medial amygdalar nucleus (MEA) exhibited heterogenicity among its different subdivisions. Its anterodorsal part (MEAad) received very dense (6+) to dense (5+) projection fibers along its rostro-caudal extent (Figure 3A-C). The anteroventral (MEAav) (Figure 3C-D), posteroventral (MEApv) (Figure 3E) and posterodorsal (MEApd) (Figure 3D-E) part were moderately (3+) labelled throughout the rostro-caudal axis. However, the boundaries of these nuclei had higher density fibers compared to the core.

#### 3.3.4 Other nuclei

The nuclei that did not fit in the above classification include the anterior amygdalar area (AAA) (Figure 3A), intercalated areas (IA) (Figure 3A-C), and posterior amygdaloid nucleus (PA) (Figure 3F), all of which were heavily innervated (*See table 2*).

**Table 2.**
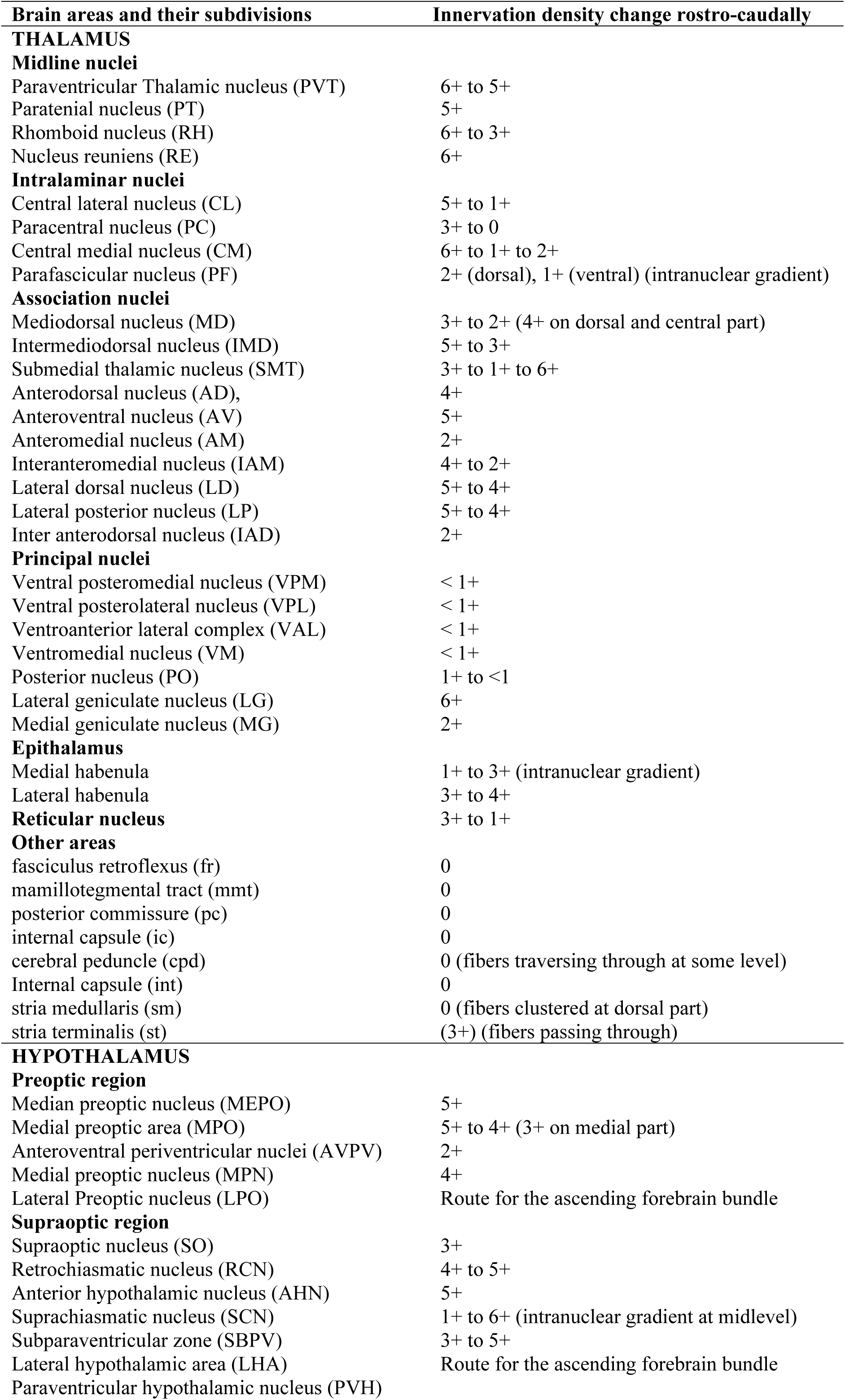
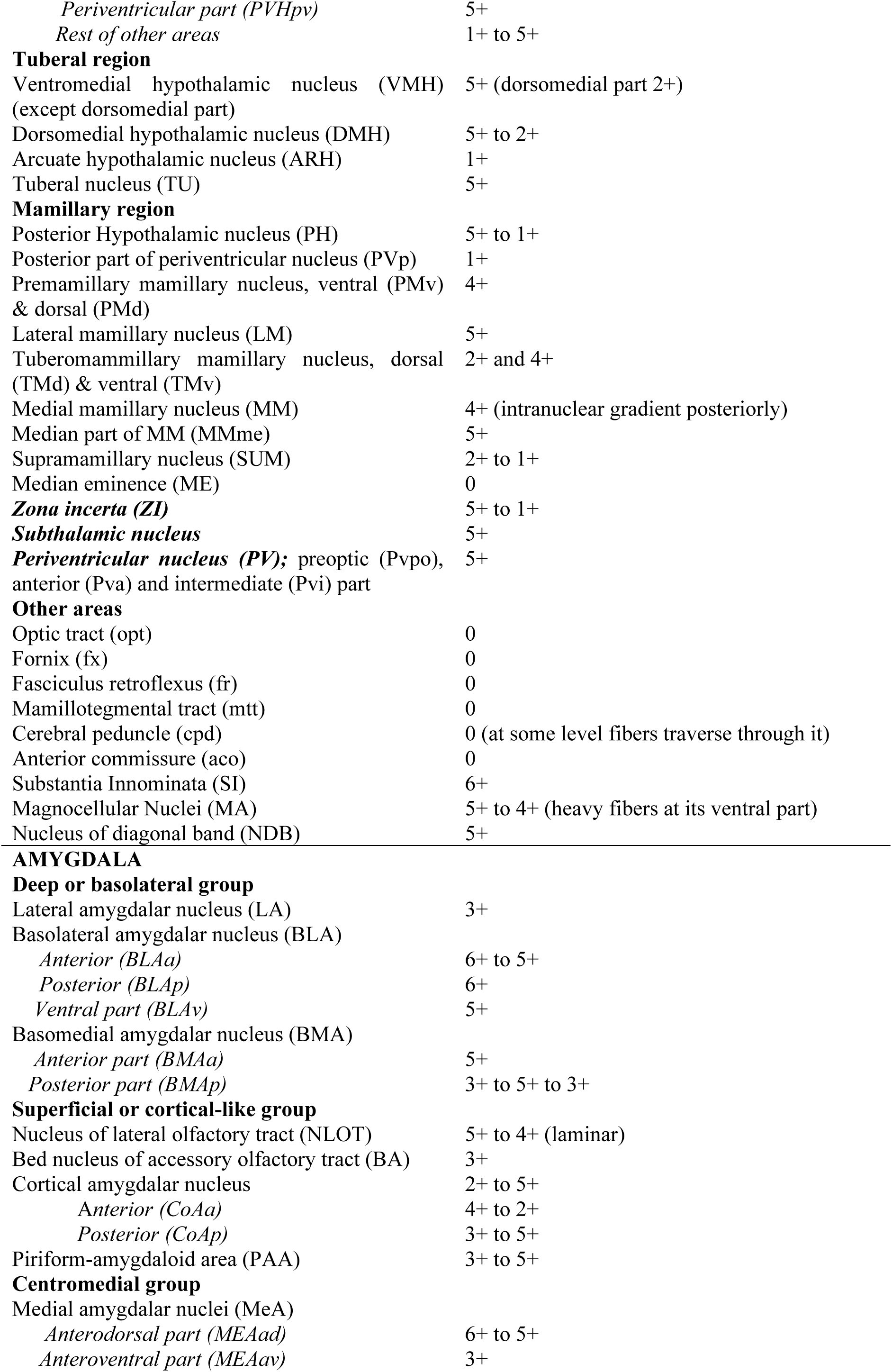
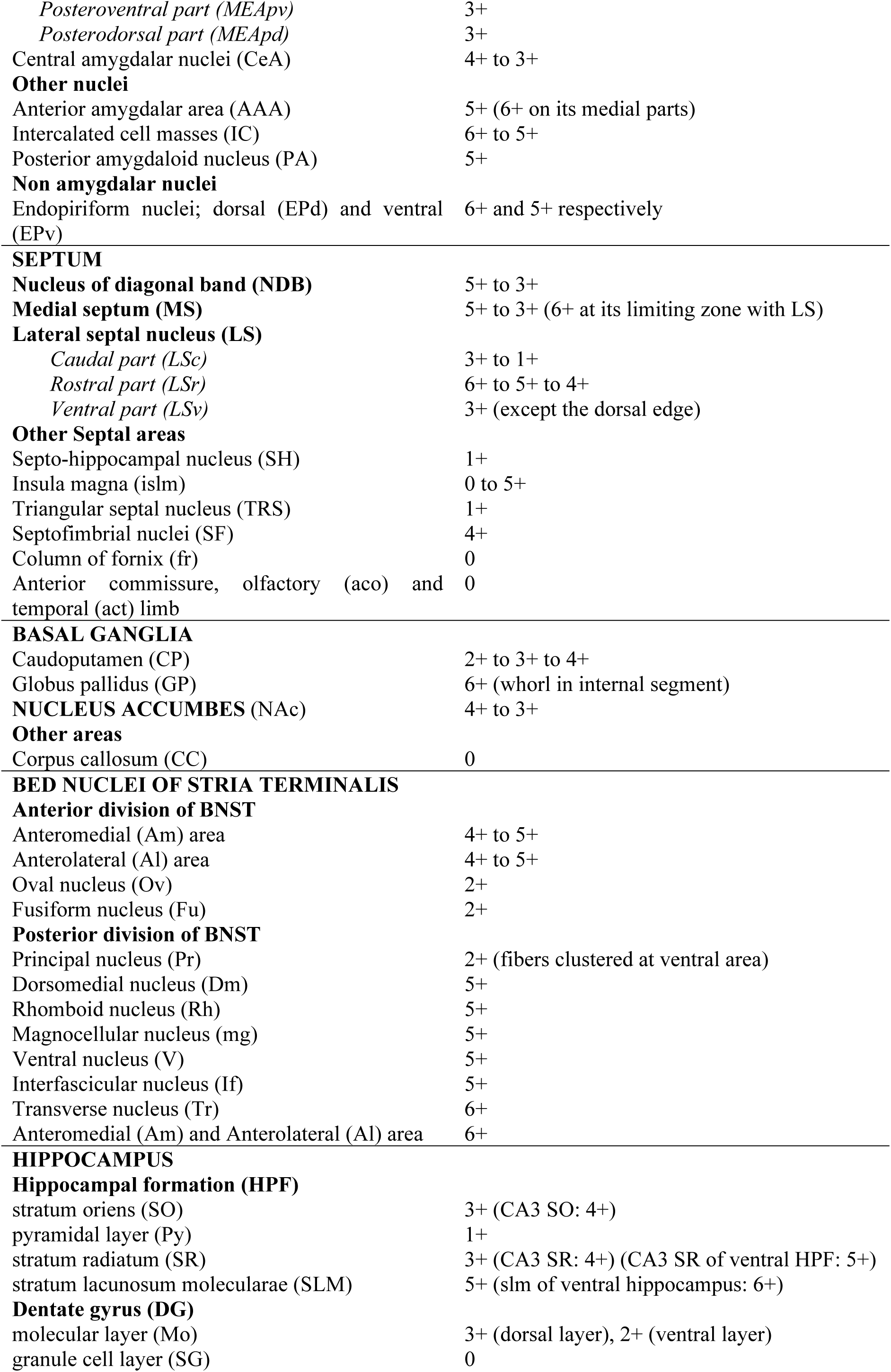
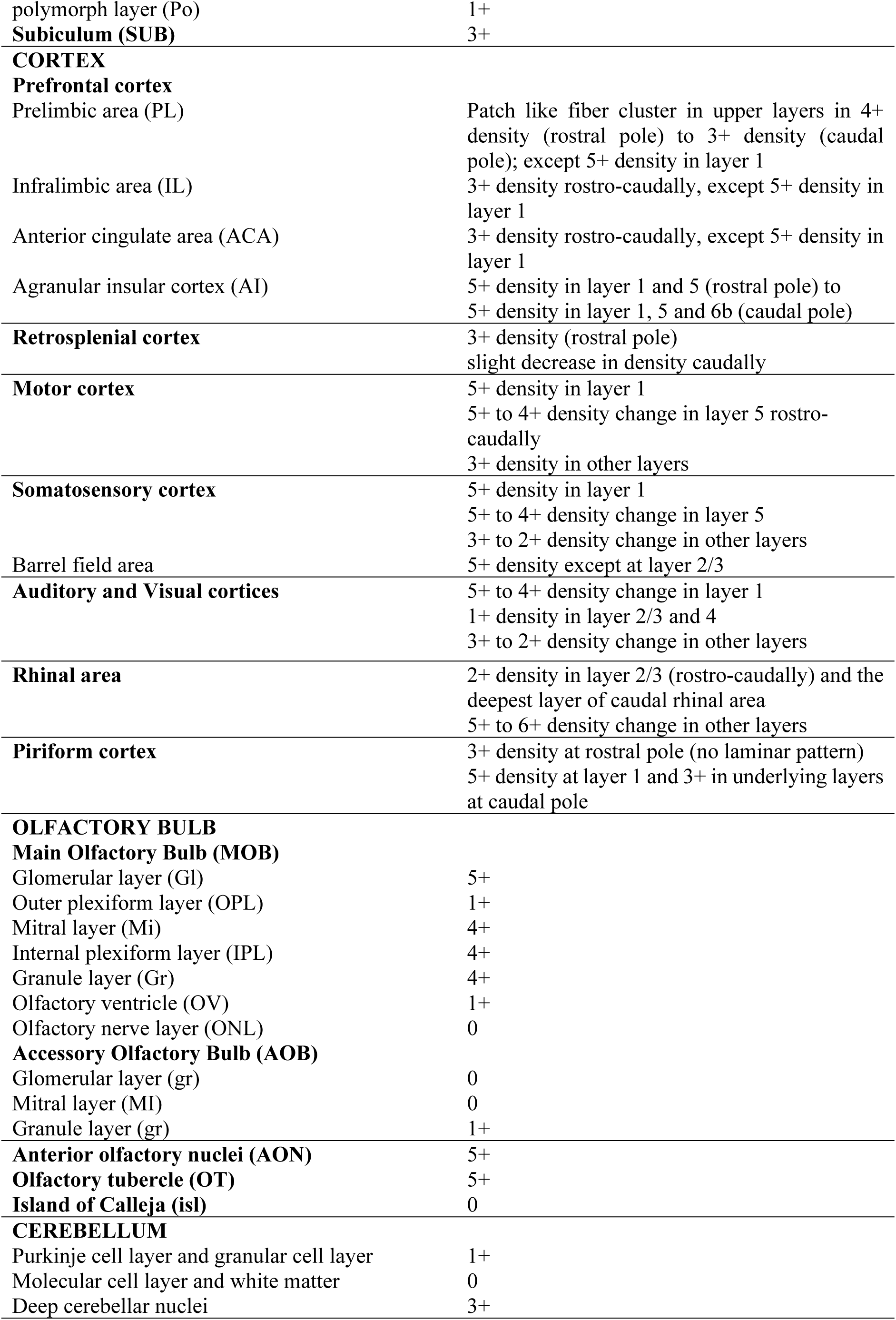
Innervation density across different brain areas and their sub-divisions

### 3.4 INNERVATION PATTERN IN THE SEPTUM

The septum has three major parts and their subdivisions. The major areas are the medial septum (MS), lateral septum (LS), and the nuclei of diagonal band (NDB) (Risold, 2004). We observed that highly fluorescent and thick collaterals of pathway fibers entered into the septum via NDB (Figure 4A), substantia innominata (SI) (Figure 5A) and via lateral preoptic area (LPO) (Figure 4E), which made the septum appear very rich in innervation. The septum served as the route for these fibers to reach above the corpus callosum (CC) (where they clustered to form the supracallosal bundle) (Figure 4A, arrow head) and into dorsal fornix (df) and fimbria (fi) (Figure 7A & E, arrow). Finally, all these fibers culminated into the hippocampus (Figure 7E). On a close inspection of the septal component, we noticed that few thin and varicosed fibers of innervation were scattered among the thick collaterals of pathway fibers. Moreover, we observed heterogenicity in the innervation density and orientation of fibers among the different nuclei.

**Figure 4:**
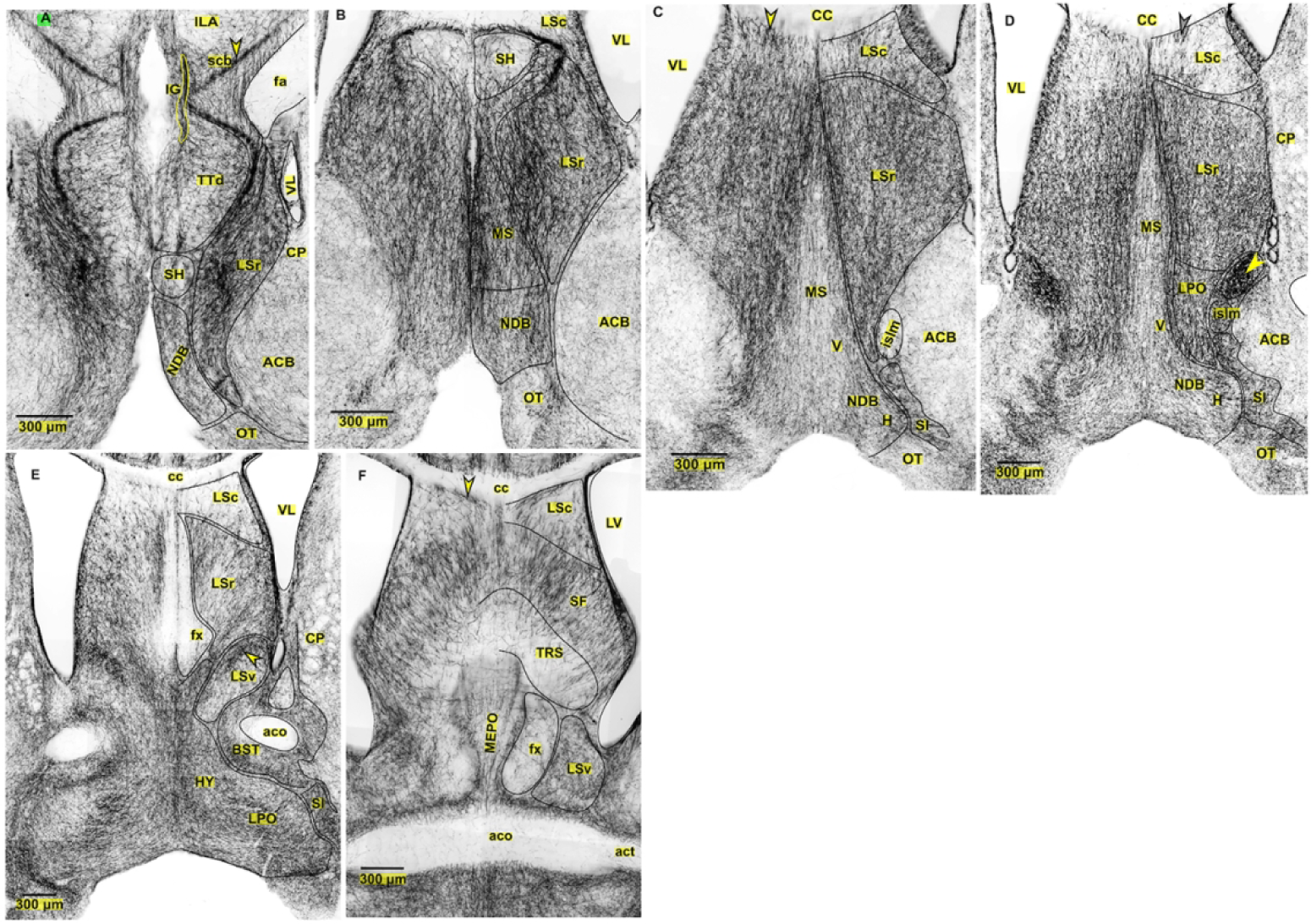
Innervation pattern of SERT-EGFP fibers across the rostro-caudal extent of septal nuclei. **(A-B)** *Sections from rostral septal level* **(A)** LSr nuclei (6+), collaterals of pathway fibers travelled through LSr to form the supracallosal bundle (scb) located above the CC (arrow head). NDB (4+). SH nuclei (2+). IG contains the collaterals of pathway fibers which innervate the medial cortex. Non-septal nuclei: TTd (3+) **(B)** NDB, MS, and LSr nuclei (5+). Ventrodorsally running collaterals of pathway fibers were clustered very densely (6+) in the medial and lateral border of MS which terminated by bounding the SH nuclei dorsally. SH nuclei (1+). **(C-D)** *Sections from mid septal level.* LSr nuclei (5+). NDB and medial part of MS (3+). Fibers can be observed entering into the septum via SI. Ventrodorsally running collaterals of pathway fibers clustered very densely (6+) in the lateral border of MS which reached upto the ventral border of CC. **(C)** LSc (3+) with vertically oriented fibers abutting the CC (arrow head). Islm **(D)** LSc (2+) with vertical fibers abutting CC (black arrow head). Very dense (6+) cluster of fibers between LSr and NAc (yellow arrow head). Islm (5+). **(E-F)** *Sections from caudal septal level.* **(E)** Medially diverted collaterals of ascending fibers from LPO nuclei entered into the septum. LSr nuclei (4+). Mediolaterally, directed fibers in LSr. LSv nuclei (3+), except densely clustered fibers at its dorsal part (arrow). LSc (1+) with vertical fibers at the dorsal border. Column of fornix (fx) (0). Thin fascicle of fiber passing ventrodorsally in between the two columns. **(F)** SF nuclei (4+); mediolaterally oriented fibers. LSc nuclei (1+) with vertical fibers at its dorsal border (arrow head). TRS (1+). LSv nucleus (3+).

**Figure 5:**
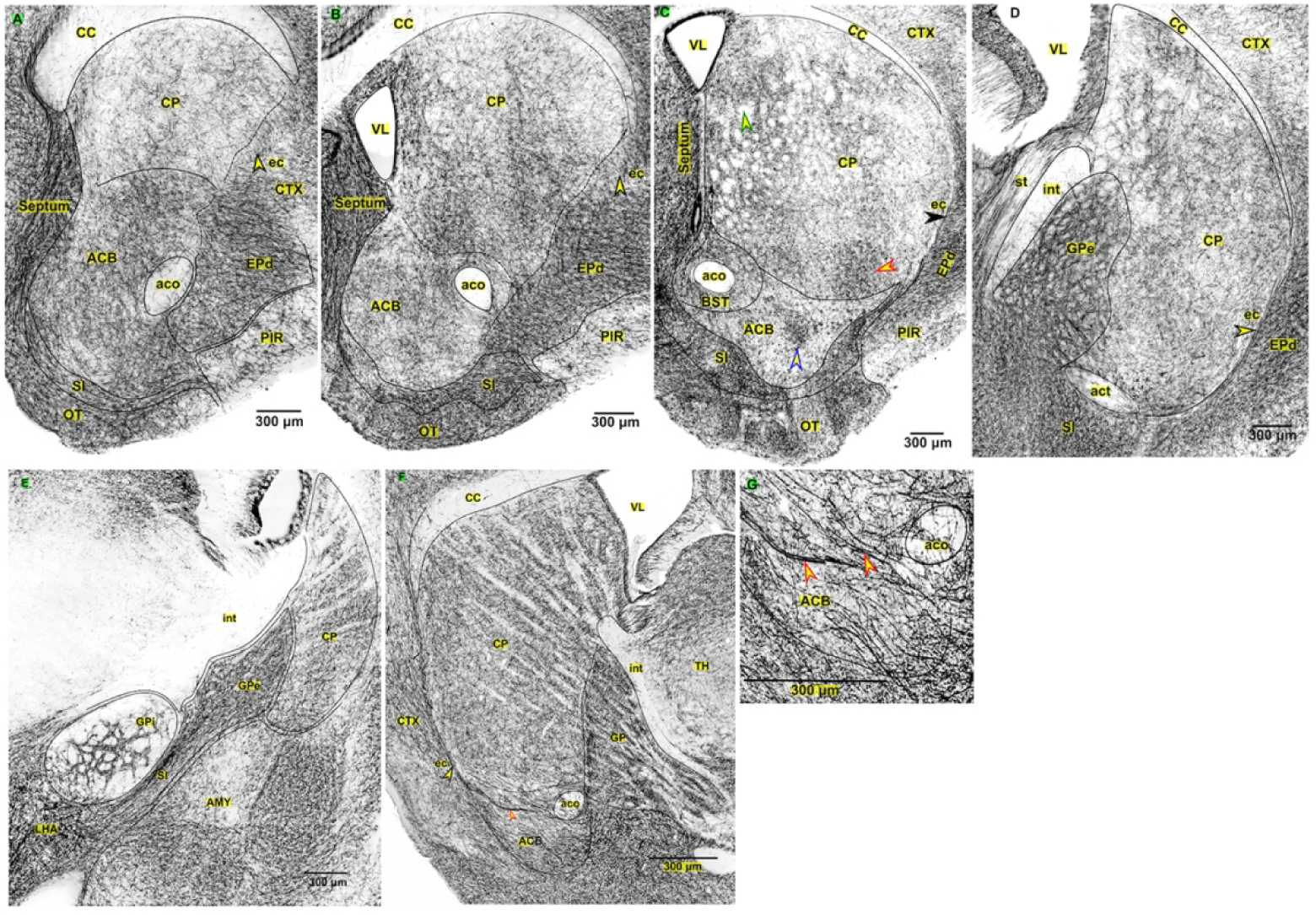
Innervation pattern of SERT-EGFP fibers across the rostro-caudal extent of caudoputamen and nucleus accumbens. *(The images are the serial sections arranged rostro-caudally)* **(A)** CP (2+). NAc (4+). Fibers entered into the CP and NAc through collaterals of fibers running in external capsule (arrow head) (A-C). **(B)** CP (3+ on lateral area and 2+ on medio-dorsal part). Fibers running within external capsule (arrow head) wind up around olfactory limb of anterior commissure (aco) or directly enter into the CP and NAc (3+). **(C)** collaterals of pathway fibers within SI wind around aco to enter into the medial side of CP (segmented black arrow line). Medial part (3+) have higher innervation compared to lateral (2+). Region of thalamocortical fibers traversing through CP appear as circular gap (green arrow head). Collaterals of ascending pathway fibers appear as patch in CP (red arrow head) and NAc (blue arrow head). NAc (2+), in the area except the patch (blue arrow head) **(D)** CP (3+ homogeneous distribution throughout). GPe (6+). Empty holes in the CP and GPe are areas of non-labelled thalamocortical fibers. Collaterals of pathway fibers in the SI, directly innervated the GPe. **(E)** CP (4+). GPe (6+). Fibers arising from main ascending bundle at LHA entering into the GPe through SI. Fibers were very densely clustered only around the thalamocortical fibers (whorl like pattern) in the GPi and sparse elsewhere. **(F)** *Sagittal section showing both caudoputamen and nucleus accumbens.* Gap indicates unlabeled thalamocortical fibers. Thin, relatively straight fibers were seen terminating mainly in the NAc (red arrow head) and few in the CP. Some of them entered into the external capsule (yellow arrow head) which provided the innervation both to the cortex and CP. **(G)** enlarged view of nucleus NAc showing the straight, non-varicose terminal fibers seen in E (red arrow head)

**Figure 6:**
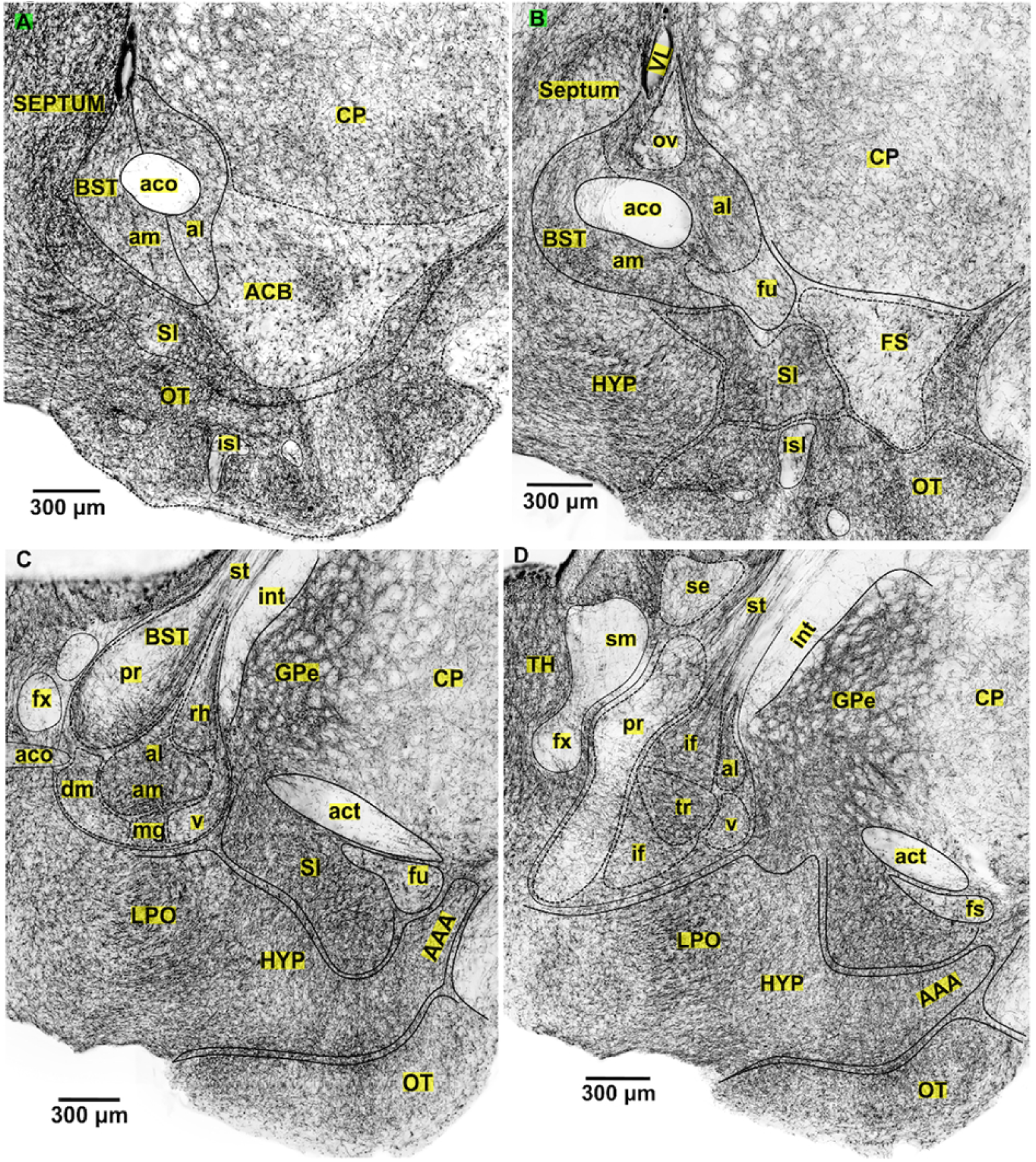
Innervation pattern of SERT-EGFP fibers across the rostro-caudal extent of bed nuclei of stria terminalis and olfactory tubercle. *(The images are the serial sections arranged rostro-caudally)* **A-B:** *anterior BNST*. Fibers entered into the BNST via collaterals of pathway fibers clustered within SI. **(A)** am and al nuclei (4+). **(B)** am and al nuclei (5+). ov and fu nuclei (2+). **C-D:** *posterior BNST*. **(C)** am and al nuclei (6+). dm, mg, v, and rh nuclei (5+). Pr nuclei (2+), except at ventral area. Fibers on pr, am, al and rh nuclei entered into the stria terminalis (st). **(D)** tr nuclei (6+). if, v, al nuclei (5+). pr nuclei (2+). Fibers from BNST entered into the stria terminalis (st) to run dorsally above the thalamus. (**A-B)** OT (6+). Fibers in OT were clustered around isl; while latter were almost devoid of labelling. (**C-D)** OT (5+); fibers distributed were fine in morphology and homogeneously distributed throughout.

**Figure 7:**
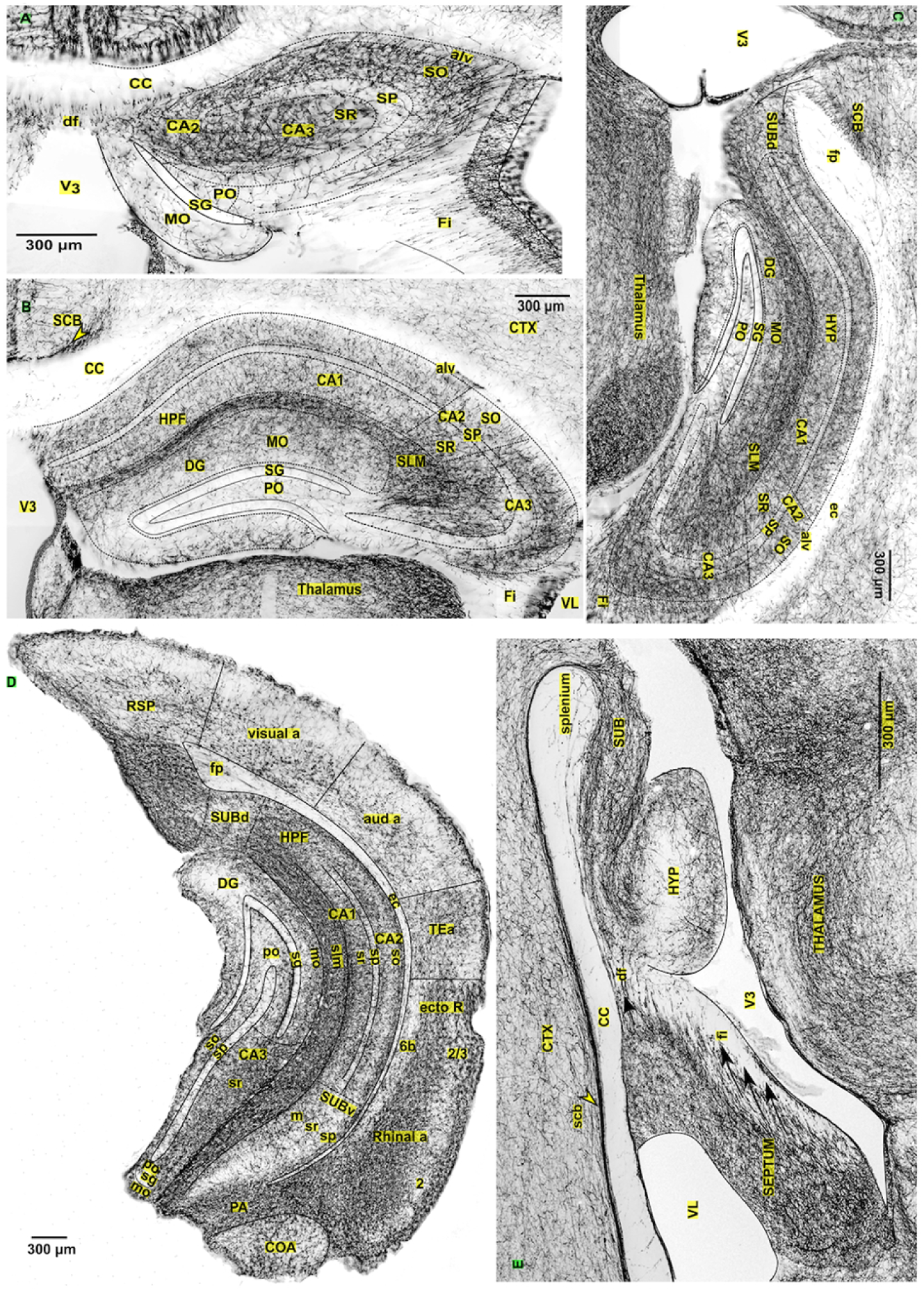
Innervation pattern of SERT-EGFP fibers across the rostro-caudal extent of hippocampus. **(A)** *Rostral end of the dorsal hippocampus*. Fibers entered into the hippocampus through dorsal fornix (df) and fimbria (fi). Some of these collaterals passed into the alveus (alv) and ramified into the SO. SO and SR (4+). SP, PO and MO layer (1+). SG (0). **(B-C)** SO and SR of CA1 and CA2 zone of HPF (3+). SO and SR of CA3 zone of HPF (4+). Whole extent of SLM (5+). Whole extent of SP (1+). Dorsal part of MO layer of DG (3+). Ventral part of MO layer of DG and PO (1+). SG (0). Fibers enter into the hippocampus through fimbria (fi) and fibers spreading laterally from SLM made the CA3 SO and CA3 SR appear more clustered with fibers respectively. **(C)** *Retrosplenial part of dorsal hypothalamus*. Collaterals of pathway fiber from supracallosal bundle (scb) passed ventrally into the hippocampus (black arrow) mainly into the alvelus (alv) and SLM. **(D)\*** *Ventral hippocampus*. SO, SR of CA1 (3+). SLM (6+). MO layer (3+) except at apex. SR of CA3 (5+). SO of CA3 (3+). SP and PO (1+). SG (0). SUBv and SUBd (3+). Fibers passing through the SLM moved out laterally to innervate the rhinal cortex. **(E)** *Image showing the sources of innervation fibers of the hippocampus.* supracallosal bundle (scb) running dorsally above corpus callosum (CC) wind around the splenium to enter the hippocampus (yellow arrow head). Collaterals of pathway fibers enter into the septal nuclei run dorso-caudally to pass into the dorsal fornix (df) and fimbria (black arrow) which subsequently innervate the hippocampus. **\*(D) *Image showing both the ventral hippocampus and cortical section from occipital pole*.** *RSP, Visual and auditory cortex*: layer 1 (4+), layer 2/3 and 4 (1+), underlying layers (2+). *Temporal association area (TEa)* (3+). *Rhinal cortex*: 6+ except in second and deepest layer.

#### 3.4.1 The nucleus of diagonal band (NBD) and medial septum (MS)

The NDB consists of two limbs, the vertical and horizontal limb (Figure 4C, D). The vertical limb is in continuation with the medial septum (MS) and occupies the medial most position in the rostral half of the septum (Figure 4B-D). Towards the rostral pole, we observed that the medially diverted collaterals of ascending pathway fibers ran dorsally along the lateral border of NDB and via the lateral septum (LSr) to reach above the CC, thus forming the supracallosal bundle (SCB) (Figure 4A). The SCB ran caudally over the CC while providing the innervation to the medial part of cortex en-route which finally turned around the splenium of CC to enter into the hippocampus (Figure 7E). In the slight caudal section, some of the medially diverted collaterals of ascending fiber ran dorsally along lateral border of NDB to continue into the MS which then terminated upon surrounding the septo-hippocampal (SH) nuclei (Figure 4B). We observed that at the midway septal level, majority of the fibers entering into the septum along the lateral border of NDB appeared relatively straight and oriented ventro-dorsally to reach up to the ventral border of CC (Figure 4C-D). They gave off many projections to the lateral septum en-route. A particularly striking feature of NDB and MS of the mid-level septum was the moderately (3+) innervated zone lying close to the midline (Figure 4C-D). This zone consisted of mainly the thin fibers of innervation with very few collaterals of ascending pathway fibers. The overall density and pattern of SERT-EGFP fibers in the MS were similar to that of NDB. In addition, we observed a strikingly very dense (6+) cluster of fibers in the limiting zone between nucleus accumbens (NAc) and lateral septum (LSr) (Figure 4D, yellow arrow head).

#### 3.4.2 Lateral septal nucleus

The lateral septum (LS) consists of three main subdivisions; the caudal (LSc), rostral (LSr) and ventral (LSv). We noticed heterogeneous distribution of SERT-EGFP fibers among the different subdivisions of the LS. The LSc was the least innervated area of the whole septum. It contained vertically oriented fibers at its border abutting the CC (Figure 4C-F, arrow head). Upon tracing these vertical fibers, we noticed they passed caudally into the dorsal fornix which served as one of the sources of innervation of the hippocampus (Figure 7A). The rest of the area of LSc had randomly oriented fibers in minimal density (1+) (Figure 4D-F). However, LSc was moderately labelled at its rostral pole (Figure 4C). Similarly, we found that LSv was moderately (3+) innervated (Figure 4E-F). However, the collaterals of ascending fiber entering into the septum were arborized heavily on its dorsomedial part (arrow head, Figure 4E). We observed that the LSr was among the densely labelled areas of the whole brain, however, it demonstrated heterogenicity in the labelling density and orientation of fibers along the rostro-caudal axis. It was very densely (6+) innervated in the rostral half (Figure 4A-B), densely (5+) with randomly oriented fibers at the mid-level (Figure 4C-D) and the density decreased even further towards its caudal pole (4+) where the fibers were mediolaterally oriented (Figure 4E). Upon tracing the fibers, we noticed that the collaterals of pathway fibers within the LSr passed caudally into septofimbrial nuclei (SF) (Figure 4F) and finally were projected into the fimbria. The fimbria (fi) served as one of the sources of innervation of hippocampus (Figure 7A & E). Thus, the medially diverted fibers from the main ascending bundle traversed across the different level of septum to reach the supracallosal bundle, fimbria and dorsal fornix which finally culminated into the hippocampus (Figure 7E).

#### 3.4.3 Other Septal areas

There are other areas which are located within the septum but are not its principal components. We observed that septo-hippocampal (SH) nucleus was sparsely (1+) innervated (Figure 4A-B). However, the collaterals of pathway fibers running dorsally across MS terminated along its outskirts (Figure 4B). Innervation in the insula magna (islm), the largest islands of Calleja, exhibited the rostro-caudal gradient. The rostral *islm* that was almost devoid of the innervation (Figure 4C) received dense (5+) network of fibers at its caudal extent (Figure 4D). Similarly, column of fornix (fx) appeared completely devoid of the innervation except a thin fascicle of fibers that was observed running dorsally in-between two bilateral columns which reached up to the ventral border of CC (Figure 4E). Triangular septal nucleus (TRS) was sparsely (1+) innervated (Figure 4F). However, SF contained mediolaterally oriented collaterals of pathway fibers in moderately high (4+) density (Figure 4F).

### 3.5 INNERVATION PATTERN IN THE BASAL GANGLIA

We observed gradual increase in the labelling density of caudo-putamen (CP) rostro-caudally. The mildly (2+) labelled CP towards its rostral extent (Figure 5A) received moderately high (4+) projections towards its caudal pole (Figure 5E). We noticed that the collaterals of main ascending bundle clustered in SI were diverted either medially or laterally to provide the innervation to the CP (Figure 5A-D). The laterally directed fibers passed along external capsule (ec) (Figure 5A, arrow head) and/or turned around the anterior commissure (aco) to innervate the lateral part of CP (Figure 5B), whereas the medially directed collaterals of pathway fibers passed dorsally across BNST around the *aco* to reach the medial side of CP (Figure 5C, segmented black arrow line). Thus, because of these different routes of innervation, the density of labelling in the CP was higher either laterally (Figure 5B) or medially (Figure 5C) at different levels along the rostro-caudal extent. In addition, the collaterals of pathway fibers running rostrally through CP were clustered together at some points which appeared as patches in the coronal section (arrow Figure 5C, red arrow head). We noticed that the areas through which thalamocortical fibers were traversing ahead appeared as circular gaps in the coronal sections as they were unlabeled with GFP (Figure 5C, green arrow head). The SERT-EGFP fibers surrounding these gaps gave it a whorl like appearance. In sagittal sections, the collaterals of pathway fibers entering into the CP via external capsule or GP can be distinguished as smooth, straight, large diameter structures which were more evident in the NAc (Figure 5F & G, red arrow head). The terminal fibers innervating CP were very fine in morphology. The GP received very dense (6+) serotonergic projection which made it clearly distinguishable from the adjacently located striatum (Figure 5D-E). The collaterals of main ascending fibers streamed into GP from lateral hypothalamic area (LHA) through SI (Figure 5E). In addition, we observed that the pattern of SERT-EGFP fibers in GP external segment (GPe) changed from a whorl pattern (Figure 5D) to homogeneous labelling (Figure 5E) rostro-caudally.

### 3.6 INNERVATION PATTERN IN THE NUCLEUS ACCUMBENS

The innervation pattern in the nucleus accumbens (NAc) was unique because of the presence of two different types of terminal fibers. We observed a unique type of fine, non-varicose, relatively straight terminals scattered among the ubiquitous punctate type of fibers of innervation. These unique types of terminal fiber had morphology similar to pathway fibers but were thin in diameter. They appeared as loosely clustered patch in the coronal sections of caudal NAc (Figure 5C, blue arrow head) and can be visualized much better in the sagittal sections (Figure 5F & G, red arrow head). We noticed that the innervation density in the NAc was higher rostrally (4+) (Figure 5A) than its caudal pole (3+) (Figure 5C). This was because the loosely clustered fibers at the caudal NAc were scattered throughout the nuclei at the rostral pole.

### 3.7 INNERVATION PATTERN IN THE BED NUCLEUS OF STRIA TERMINALIS

We observed that collaterals of pathway fibers clustered within substantia innominata (SI) were diverted medially which turned around the anterior commissure (aco) to reach the medial side of CP; thus, innervating the bed nuclei of stria terminalis (BNST) en-route (Figure 6A-B and 5C). However, towards the caudal pole of BNST, the detached fibers from the main ascending bundle at lateral preoptic area (LPO) entered into the BNST which then continued into the stria terminalis (st) (Figure 6C-D). Those fibers in ST ran caudally above the thalamus to project into the amygdala ultimately (Figure 3C-D). In the anterior division of BNST, we observed that the SERT-EGFP fibers were distributed homogeneously in moderately high density (4+) to dense (5+) levels within the anteromedial (am) and anterolateral (al) nuclei. However, the density was mild (2+) in the oval (ov) and fusiform nuclei (fu) (Figure 6A-B). At the posterior division of BNST, we noticed that anteromedial (am), anterolateral (al) and transverse nuclei were very densely (6+) labelled. The rest of the nuclei had dense (5+) labeling except the principal nuclei (pr) which was mildly (2+) labelled except on its ventral part (Figure 6C, D).

### 3.8 INNERVATION PATTERN IN THE HIPPOCAMPUS

Hippocampus constitutes of 4 major parts, viz: hippocampal formation (HPF), dentate gyrus (DG), subiculum (SUB) and entorhinal area (ENT). HPF which is also known as cornu ammonis (CA) is further subdivided into various zones, viz; CA1, CA2 and CA3. HPF consists of four different layers; from outward to inward; viz: stratum oriens (SO), pyramidal layer (Py), stratum radiatum (SR), stratum lacunosum molecularae (SLM), whereas, the DG comprises of three layers viz; molecular layer (Mo), granule cell layer (SG) and polymorph layer (Po) (Witter and Amaral, 2004).

We observed heterogeneously distributed SERT-EGFP fibers among the various layers of hippocampus that gave it a laminar appearance. The SG of dentate gyrus was almost devoid of the innervation, whereas fibers were sparsely (1+) scattered within SP and PO layers (Figure 7A-C). SLM was densely (5+) labelled in dorsal hippocampus (Figure 7B-C) which became even more dense (6+) in the ventral hippocampus (Figure 7D). The compactly arranged fibers within CA1 SLM were diffused laterally to distribute over the CA3 SR in moderately high (4+) density (Figure 7B-C). Similarly, the fibers entering into the hippocampus via fimbria (fi) also labelled the CA3 SO in moderately high (4+) density. Therefore, except the CA3 SR and CA3 SO, the rest of the areas of these layers and the dorsal part of MO layer were moderately (3+) labelled, whereas the SERT-EGFP fibers were mildly scattered in the ventral part of MO layer (Figure 7B-C). The pattern in the ventral hippocampus was almost congruent with dorsal hippocampus (Figure 7D) except the labelling in SLM was higher than its dorsal counterpart.

We traced the fibers of innervation to the hippocampus which were derived through three different sources, viz; dorsal fornix (df), supracallosal bundle (scb) and fimbria (fi). The medially diverted collaterals from the main ascending bundle that traversed through septal nuclei were finally streamed into the scb, df and fi (Figure 7E) (also see the innervation to the septum above). The dorsal fornix was mainly responsible for the innervation of rostral pole of dorsal hippocampus (Figure 7A). The fibers from the fimbria projected into the hippocampus through laterally located CA3 zone (Figure 7A-C). The supracallosal fiber bundle ran caudally above the CC and turned around its splenium to terminate finally into the hippocampus (Figure 7C black arrow & E yellow arrow head). These fibers mainly streamed into the SLM and alveus and were subsequently distributed into the adjacent layers (Figure 7C). We noticed that the collaterals of pathway fibers within the SLM of ventral hippocampus passed externally to get heavily distributed over the rhinal areas (Figure 7D).

### 3.9 INNERVATION PATTERN IN THE CORTEX

We traced various routes of innervation of the cerebral cortex which varied depending on the brain levels. At the frontal pole of the brain, the collaterals of pathway fibers clustered within the substantia innominata (SI) passed laterally around the rhinal fissure (RF) to project mainly into cortical layer 1 (Figure 8B-C). Similarly, the fibers clustered within the endopiriform nuclei (EP) also streamed into the layer 1, 5 and 6b of lateral cortex (Figure 8C-D). The collateral pathway fibers running in the external capsule (ec) supplemented the innervation of layer 6b and additionally innervated the lateral part of CP (Figure 8C-D, 5B-C arrow). The fibers were very densely (6+) distributed within the claustrum (CLA) (Figure 8C-D). We observed that the medial cortex had separate sources of innervation. It was innervated mainly by the collaterals of pathway fibers streaming through supracallosal bundle (scb) (Figure 7E, 8D & E), indusium griseum (ig) (Figure 8C & H, arrow), and medial cortical bundle (Figure 8C). We noticed that the outermost cortical layer was the most labelled (5+) layer having transversely oriented fibers to the horizontal axis. We observed difference in the innervation pattern among the various cortical areas and gradual decrease in the labelling density as traced rostro-caudally.

**Figure 8:**
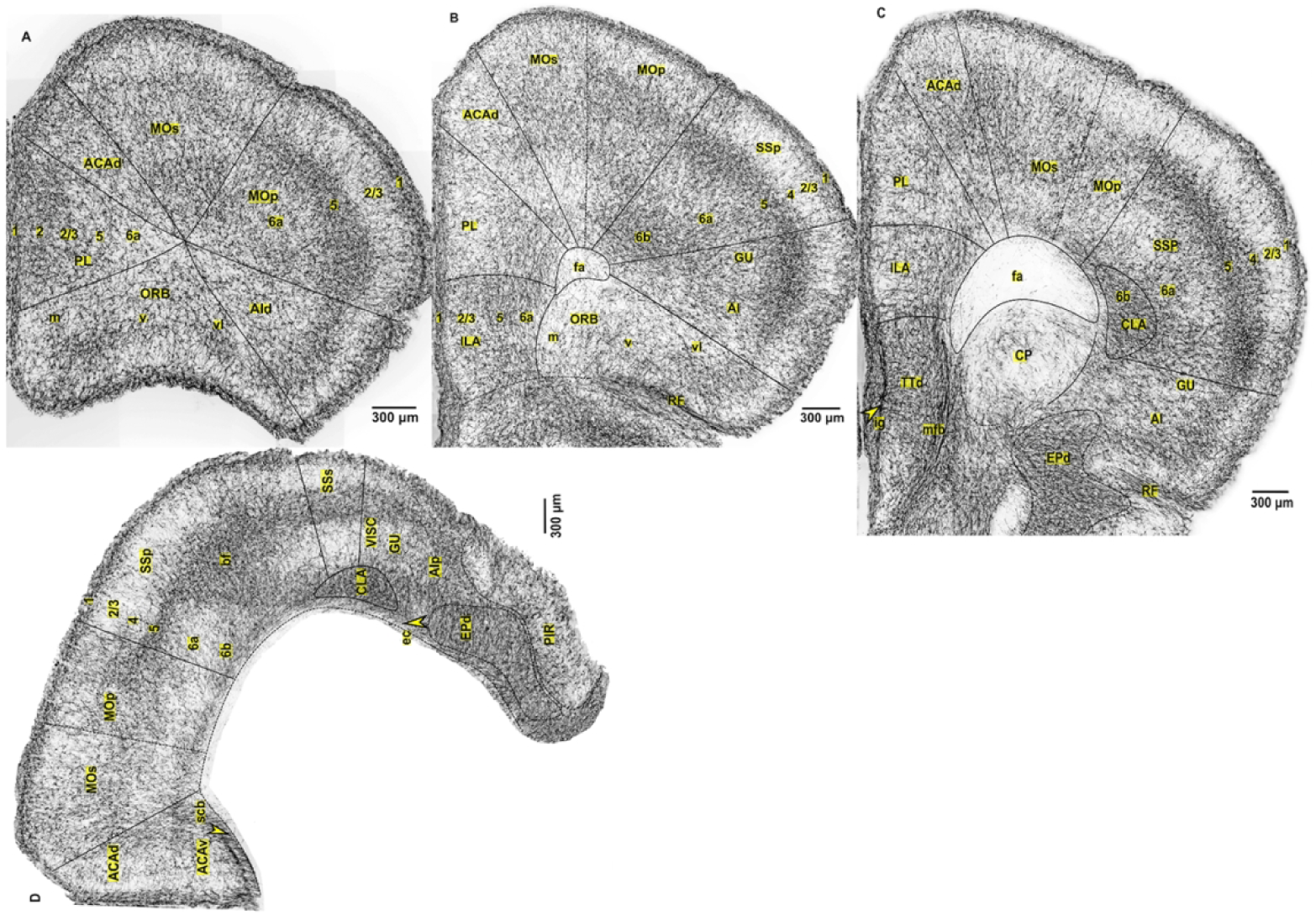
Innervation pattern of SERT-EGFP fibers across the rostro-caudal extent of cortex. *(The images are serial cortical sections arranged rostro-caudally)* **(A)** *Rostral pole of the cortex*. Layer 1 in all the areas (5+). *PL area:* patch like cluster of fibers. layer 2 and 2/3 (4+). rest of the layers (3+). *ACAd area* (3+). *MOs area:* layer (5+) and deeper part of 6a (4+), other layers (3+). *MOp area and AId area:* layer 5 (5+), rest of the layers (3+). *ORB area:* layer 5 (4+) and rest of the layers (3+) **(B)** Layer 1 in all areas (5+). *ILA, PL, and ACAd area* (3+). Fibers from endopiriform nuclei (EPd) enter into the medial cortex through the deeper layer of ILA and PL cortex. *MOs area:* 3+ in rest of the layers except 6b (2+). *MOp area:* Layer 5 (4+). Rest of the layers (3+). *SSp, GU and AI area:* layer 5 and 6b (5+). Layer 6a (3+). Layer 2/3 (2+). *ORB area:* 2+ in rest of the layers except layer 5 (3+). Fibers enter into the layer 1 of lateral cortex through areas around rhinal fissure. **(C)** Layer 1 in all areas (5+). *ILA, PL and ACAd area* (3+) in rest of the layers. Fibers enter into this medial cortex via medial forebrain bundle (mfb) (black arrow line) and indusium griseum (ig) (yellow arrow head). *MOs and Mop area:* layer 5 and 6a (3+). 2+ in rest of the layers. *SSp area:* Layer 5 and 6b (5+). Fibers clustered within the claustrum (CLA) distribute to layer 6a (3+). Layer 2/3 and 4 (2+). *GU and AI area:* layer 5 (4+). Rest of the layers (3+). Fibers clustered within EPd enter into the lateral cortex mainly through the layer 1 and Layer 5. **(D)** Layer 1 in all areas (5+). *ACAv and ACAd* (2+) except at the deeper layers where fibers from supracallosal bundle (scb, arrow head) pass dorsally towards the motor areas and horizontally into layer 6b. *MOs and MOp* area (3+) in upper layers; 2+ in deeper layers 6a and 6b. *SS area:* layer 5 (5+). Layer 6a and 6b in the barrel field (bf) area (4+). Rest of layer 6 including layer 2/3 (2+). Barrel field area in the SS cortex receive higher projections compared to rest of the layers. *Visceral area (VISC), GU and AIp area:* Fibers from EPd traverse into the lateral cortex through the layer 1, 5 and 6b (5+ in all of these layers) of these areas. Fibers travelling through external capsule (ec, arrow head) also provide innervation to deeper cortical layers and caudoputamen. *Piriform cortex (PIR):* 3+ in all of the layers without laminar pattern. **(E)** Layer 1 in all areas (5+). *RSP area* (3+). A vertical band of fibers between layer 2 and 2/3. Fibers from supracallosal bundle (scb, arrow head) pass dorsally towards the motor areas. *Motor areas (MOs and MOp):* layer 5 (4+) and rest of the layers (3+). *Somatosensory area (SSs and SSp):* layer 5 (4+) and rest of the layers (2+). **(F)** Layer 1 in all areas (5+). *Retrosplenial area:* RSPagl (3+). RSPv and RSPd (2+). *Visual area:* deeper layers (6a, 6b) (3+). Upper layers (1+). *Auditory area:* layer 5 and below (4+). Layer 2/3 and 4 (1+). *Rhinal and temporal association area (TEa):* 5+ in rest of the layers except layer 2/3 (2+). *Piriform cortex (PIR):* laminar pattern, layer 1 (5+), deeper layers (3+). **(G)** Layer 1 in all areas (5+). *Retrosplenial cortex (RSP)* (2+). *Visual and auditory area* (1+). *Rhinal and temporal association area (TEa):* 5+ in rest of the layer except layer 2/3**. (H)** *Sagittal cortical section* showing the dense cluster of collateral pathway fibers in the induseum griseum (IG) (arrow head) which is one of the main sources of innervation to the medial cortex.

#### 3.9.1 Prefrontal Cortex

The prefrontal cortex (PFC) of the mouse consists of three major parts, the medial (mPFC), orbital (ORB) and agranular insular cortices (AI). mPFC is further subdivided into prelimbic (PL), infralimbic (IL) and anterior cingulate area (ACA) (Allen Institute for Brain Science, 2004). We observed that these different areas exhibited heterogeneous innervation density and pattern along the rostro-caudal axis. At the rostral pole, the PL area had patch like fibers clustered within the upper layers in moderately high (4+) density (Figure 8A). However, the caudal extent of PL area (Figure 8C) and the whole rostro-caudal extent of ILA and ACA (Figure 8A-D) were moderately (3+) labelled except at layer 1 (Figure 8B-D). The fibers that entered into the mPFC via EP nuclei (Figure 8B) and medial cortical bundle (Figure 8C) ran vertically through its deeper layers. Similarly, the orbitofrontal cortex (ORB) exhibited an alternating laminar pattern towards its rostral extent (Figure 8A) which was less evident caudally especially on its medial (m) part (Figure 8B). Innervation of AI also appeared in a laminar pattern because of the readily identifiable layer 1 and 5 which received higher innervation compared to the intervening layer (Figure 8A-C). Traced more caudally, the deepest layer (layer 6b) in AI cortex was also densely (5+) labelled because of fibers entering into the lateral cortex through layer 1, 5 and 6b (Figure 8D).

#### 3.9.2 Retrosplenial cortex

The retrosplenial cortex (RSP) contained a distinct vertical band of fibers running in parallel to layer 1 at its rostral pole (Figure 8E & F). The fibers were moderately (3+) distributed (Figure 8E) which progressively decreased in density caudally (Figure 8F-G). The fibers clustered within the supracallosal bundle (scb) was the source of innervation to the RSP cortex (Figure 8E, arrow).

#### 3.9.3 Motor cortex

Motor cortex also exhibited the laminar innervation pattern. Similar to the other cortical areas, layer 1 was distinct because of its dense (5+) innervation. Towards the rostral pole, layer 5 of the primary motor cortex (MOp) was densely (5+) labelled (Figure 8A), which gradually decreased in density (4+) as traced caudally (Figure 8B-E). The rest of the layers were moderately (3+) labelled, thus giving it a laminar appearance. The fibers in layer 2/3 were oriented vertically to the horizontal axis, whereas randomly distributed elsewhere (Figure 8D-E). The MOp and secondary motor (MOs) area exhibited slight differences in the innervation density only at its rostral pole (Figure 8A), whereas the patterns were almost similar elsewhere (Figure 8B-E).

#### 3.9.4 Somatosensory cortex

Rostral half of somatosensory (SS) cortex appeared distinct from the adjacent motor (MO) cortex because of its densely (5+) labelled layer 5 and mildly (2+) labelled layer 2/3 and 4 (Figure 8B-D). However, the innervation density of layer 5 gradually decreased to moderately high (4+) level towards its caudal extent (Figure 8E). Thus, SS cortex appeared similar to the adjacent MO cortex at the caudal level (Figure 8E). We observed that the collaterals of pathway fibers passing within the external capsule (ec) and claustrum (CLA) densely (5+) arborized in the barrel field (bf) area (Figure 8D). The higher labelling of layer 1, 5 and 6b compared to the intervening layers made the innervation pattern alternatingly laminar in SS cortex (Figure 8B-E). Basically, we found no difference in innervation pattern between the primary and secondary SS cortex. The overall innervation density in the SS cortex decreased rostro-caudally.

#### 3.9.5 Auditory and visual cortexes

The auditory and visual cortices which are located towards the caudal pole of the mouse brain received relatively less serotonergic innervation compared to the rostrally located cortical regions like the motor and somatosensory cortices (Figure 8F-G & 7D). The outer most layer 1 was thin but densely (5+) innervated. The underlying layer 2/3 and 4 were sparsely (1+) innervated (Figure 8F-G & 7D), however, the innervation density in rest of the layers decreased from moderate (3+) labeling in the rostral half (Figure 8F-G) to milder (2+) density towards the caudal end (Figure 7D). Thus, the overall innervation density in the auditory and visual cortex also progressively decreased rostro-caudally (Figure 8F-G & 7D).

#### 3.9.6 Rhinal Area

The rhinal cortex was the most labelled cortical area. At its rostral extent, all the layers except the layer 2/3 were densely (5+) innervated (Figure 8F). However, towards its caudal pole, layer 2/3 and the deepest layer were mildly (2+) innervated while the others received very dense (6+) innervation (Figure 7D). We observed that the collaterals of pathway fibers running across the stratum lacunosum molecularae (slm) of the ventral hippocampus moved outside of it to heavily innervate the laterally located rhinal cortex (Figure 7D).

#### 3.9.7 Piriform cortex

In this three-layered structure, the rostral extent of piriform cortex (PIR) was moderately (3+) labelled without laminar appearance (Figure 8D). However, a laminar pattern was observed in the caudal PIR where the outer layer was densely (5+) labelled and appeared distinct from underlying layer 2 and 3 which were moderately (3+) labelled (Figure 3C).

### 3.10 INNERVATION PATTERN IN THE OLFACTORY TUBERCLE and OLFACTORY BULB

We observed densely (5+) distributed thick and tortuous fibers clustered around the island of Calleja (isl) within the rostral extent of olfactory tubercle (OT) (Figure 6A-B), while the *isl* being almost devoid of the innervation. However, the caudal OT was innervated with homogenously distributed fine fibers in slightly less density compared to its rostral counterparts (Figure 6C-D). The OT and SI served as the route for the ascending forebrain bundle to traverse rostrally towards the olfactory bulb (Figure 6A).

Main olfactory bulb (MOB) was the terminal projection site of the ascending forebrain bundle arising from the midbrain raphe nuclei (Figure 9A). Fibers in MOB were arranged in a laminar pattern. We noticed that the outer most glomerular (GL) layer was densely (5+) innervated with SERT-EGFP fibers. Mitral (MI), internal plexiform (iPL) and granule (GR) cell layers were readily undistinguishable because of the homogeneously distributed fibers in moderately high (4+) density. The outer plexiform layer (OPL) and olfactory ventricles (OV) were sparsely (1+) labelled. We noticed that glomerular fibers were thick in diameter, intensely labelled and contained large varicosities, while thinner fibers predominated in the infraglomerular layers. The olfactory nerve layer (onl) was completely devoid of the serotonergic projections (Figure 9A-B).

**Figure 9:**
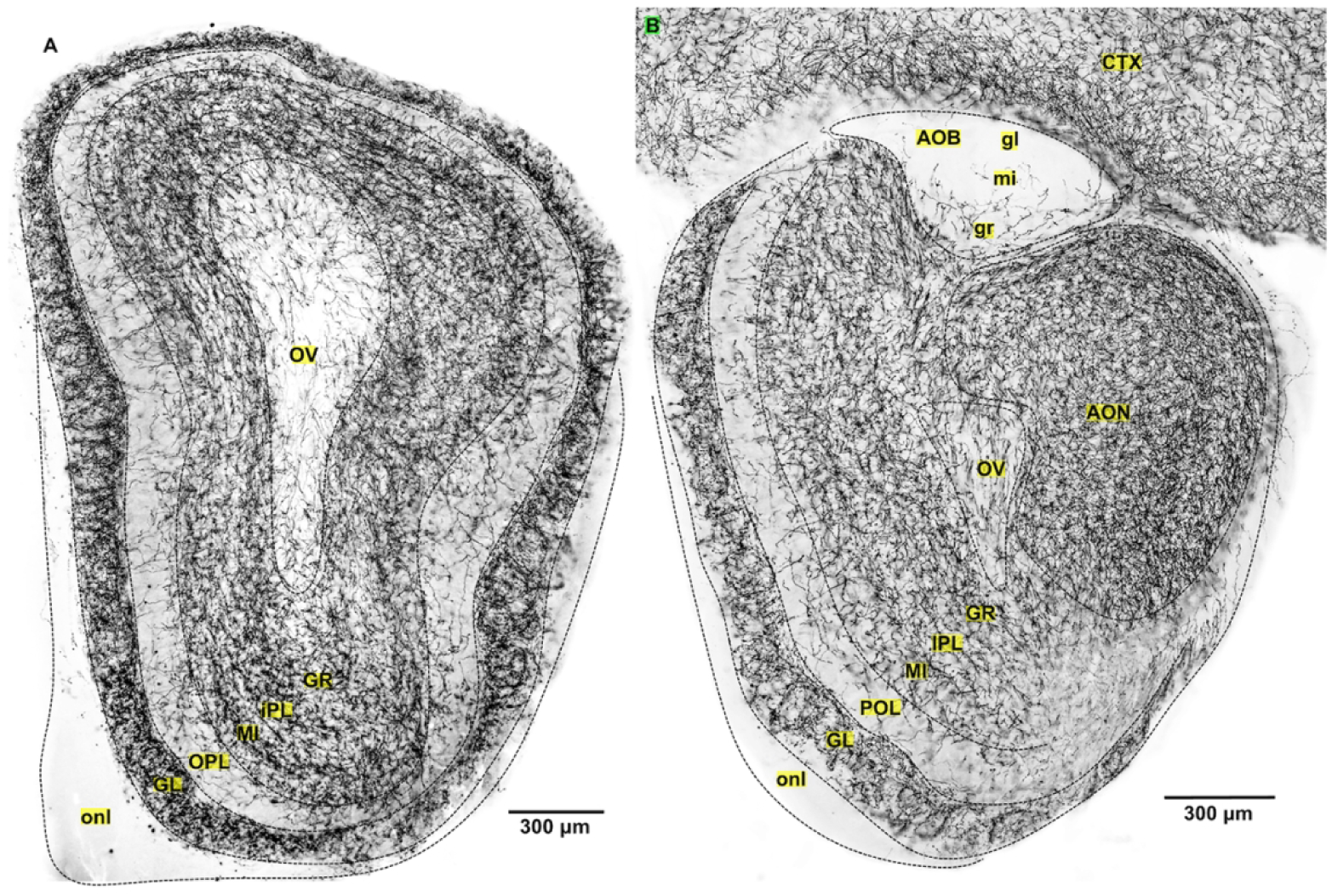
Innervation pattern of SERT-EGFP fibers in the olfactory bulb. **(A)** SERT-EGFP fibers arrangement in the main olfactory bulb (MOB) exhibit the laminar pattern. GL (5+). OPL (1+). MI, IPL and GR layer (4+). OV (1+). Optic nerve layer (onl) (0). Accessory olfactory bulb (AOB). GR layer (1+). MI and GL layer (0). Anterior olfactory nuclei (AON) acted as the route for passage for the collaterals of ascending fibers reaching to MOB

The accessory olfactory bulb (AOB) was devoid of the innervation, except the few sparsely (1+) scattered fibers within the granular (gr) layer (Figure 9B).

### 3.11 INNERVATION PATTERN IN THE CEREBELLUM

Cerebellum was the least innervated major brain component (Figure 10). Few fibers (1+) were distributed within the Purkinje cell layer (pr) and granular layer (gr). Molecular layer (mo) and white matter (wm) appeared unlabeled. Surprisingly, we observed that deep cerebellar nuclei were moderately (3+) labelled because of the direct projection from the raphe nuclei.

**Figure 10.**
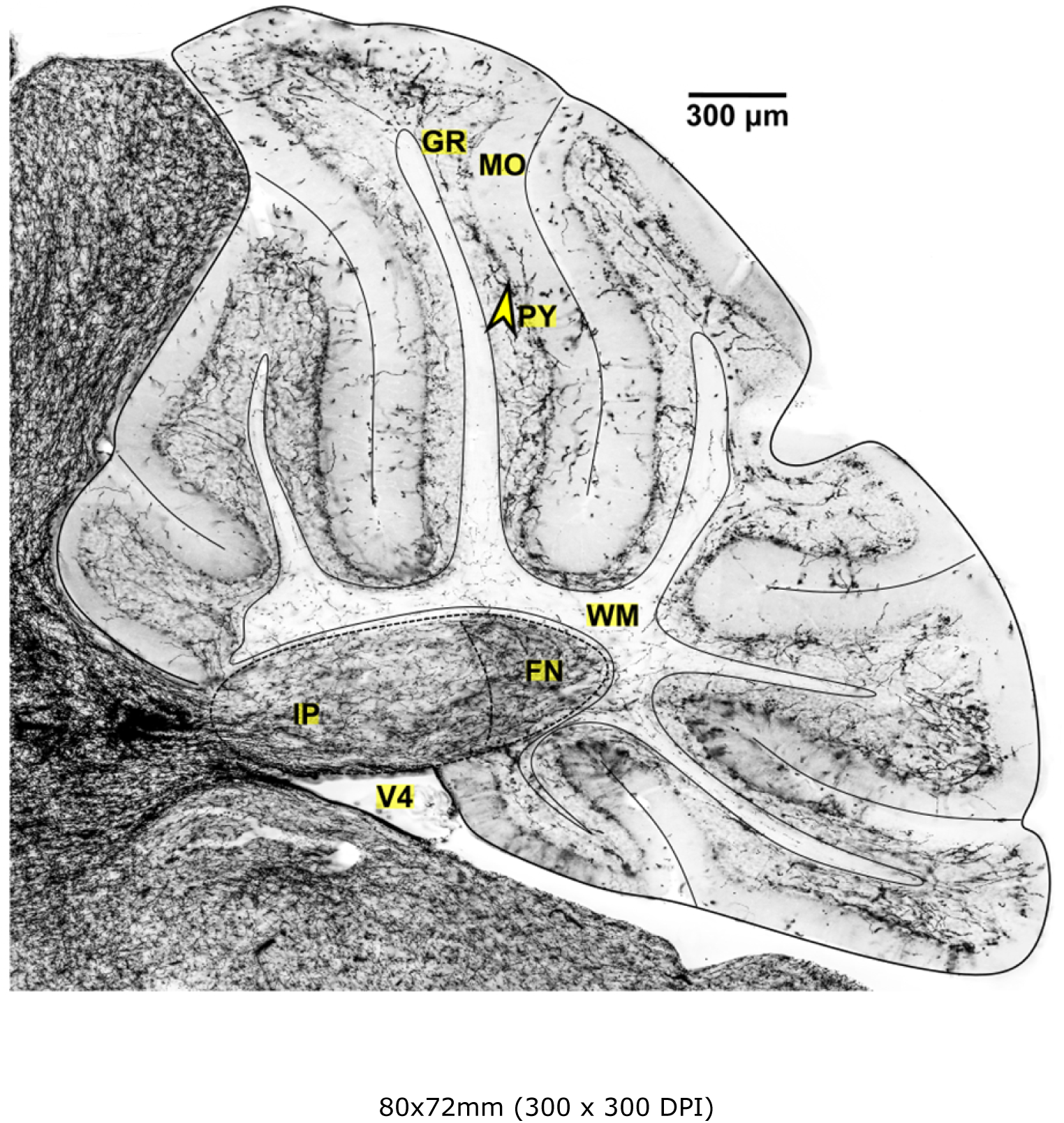
Innervation pattern of SERT-EGFP fibers in the cerebellum. Sparsely (1+) labelled layers of the cerebellum. Outermost molecular layer (MO) and white matter (WM) tree almost appear scanty. Fibers are mainly distributed in the pyramidal (py, arrow head) or granular (gr) layer. Moderate (3+) labelling within the deep cerebellar nuclei (IP and FN). Fibers from raphe nuclei passed dorsally directly into the deep nuclei.

*The results for each brain region and their nuclei are tabulated in the table 2*.

## 4. DISCUSSION

The serotonergic system is one of the diffusively organized brain neuronal systems. Although the cellular localization and projections map was first described long ago with the advent of histofluorescence technique (Dahlström and Fuxe, 1964), the improved topographical description was possible only after the development of antibodies against the enzyme tryptophan hydroxylase (Pickel et al., 1977) or the putative neurotransmitter 5-HT itself (Steinbusch, 1981). However, these techniques were accompanied with certain limitations. In order to overcome the limitations of earlier studies, we employed the genetically engineered transgenic mice expressing GFP in the 5-HT transporter. Using this mouse model, we attempted to provide more comprehensive map of the terminal field of serotonergic neurons than is currently available. In the following section, we highlight some of our results, compare them with those of previous studies and discuss the possible functional implications.

The serotonergic neurons projecting to the forebrain originated from the rostral group of raphe nuclei in the midbrain and virtually innervated all the brain regions with striking density in the hypothalamus, septal nuclei, thalamus, amygdala, olfactory bulb followed by basal nuclei and cortex. Except the white matter structures, there was hardly no brain region that did not receive 5-HT innervation. This was consistent with the previous findings (Steinbusch, 1981). We observed two different types of innervation fibers; fibers with large and spherical varicosities and fine fibers with small and granule shaped varicosities. Previous studies have shown that the difference in fiber morphology possesses functional significance. For instance, the fine 5-HT axon terminals are supposed to be extremely vulnerable to psychotropic drugs like amphetamine (O’Hearn et al., 1988). Similarly, change in the ratio of these fiber types has been observed in epilepsy where fibers with small-sized varicosities were decreased in the dentate gyrus of hippocampus, infralimbic cortex and medial septum while that of the fibers with larger-sized varicosities were increased (Maia et al., 2019). Moreover, changes in the morphology of serotonergic fibers associated with aging have been reported (Nishimura et al., 1998). Thus, analysis of 5-HT structural system could help understand the pathophysiology of mental disorders and lead to drug discovery. The 5-HT projection demonstrated an extensive and very specific innervation pattern in the brain areas as discussed below.

### 4.1 Thalamus

SERT-EGFP fibers were mainly concentrated in midline thalamus (paraventricular, paratenial, rhomboid and reuniens nuclei), rostral part of intralaminar thalamus (central medial and central lateral nuclei), some part of anterior thalamus (anteroventral, rostral part of intermediodorsal nuclei) and in the other nuclei like lateral dorsal nucleus (LD), lateral posterior nucleus (LP) and lateral geniculate (LG) complex etc.

Most of the previous findings of thalamic mapping using peroxidase-antiperoxidase (PAP) technique were consistent to ours (Cropper et al., 1984). However, they reported light labeling in several nuclei like mediodorsal, centromedian, and subthalamic nuclei where we found fibers in comparatively higher densities. Conversely, we found very light labelling in caudal part of reticular nuclei, ventral medial nuclei and posterior nuclei which they reported to be moderately labelled. Similar to us, other studies have also reported that principal thalamic nuclei lack the serotonergic input (Vertes et al., 1999). We observed very distinct distribution of fibers across the rostro-caudal axis of the reticular nuclei that progressively decreased in their density towards their caudal pole. This contrasts with the findings of one of the immunohistochemical studies which reported to have little difference (Rodríguez et al., 2011). We noticed some species differences as well. Similar to mice, the non-specific nuclei in primate (squirrel monkey) received the heaviest innervation. However, the lightly labelled reticular nuclei in mice were heavily labelled in the monkey, whereas, the richly innervated nuclei in mice such as AV, LD, and LGd were less innervated in the monkey (Lavoie and Parent, 1991).

Studies have shown that the midline thalamic nuclei in connection with limbic subcortical and cortical sites (Su and Bentivoglio, 1990) exert an arousing effect on the limbic forebrain (Vertes, 2006). The high density of 5-HT innervation in these nuclei suggests that they might modulate the emotional and cognitive functions. Similarly, the proposal of anterior thalamic nuclei as an extended component hippocampal-dependent memory network, (Aggleton and Brown, 1999) suggests that high serotonergic innervation in these nuclei might exert some effect on episodic memory. Projections of large numbers of SERT-EGFP fibers to the lateral dorsal nuclei (LD) and lateral geniculate nuclei suggest that it might modulate visually guided spatial navigation and learning (Mizumori et al., 1994) or may sharpen the visuospatial processing activity (Groenewegen and Witter, 2004). Similarly, studies have reported that the downregulation of serotonergic system in the lateral habenula is linked with the depressive symptoms in patients with Parkinson’s disease (Sourani et al., 2012). However, the precise role served by 5-HT in various thalamic nuclei and their function remains to be fully determined.

### 4.2 Hypothalamus

We observed that the hypothalamic nuclei received strong SERT-EGFP fiber input with exceptions in some nuclei. The ascending forebrain bundle traversed rostrally through the lateral hypothalamic area (LHA) and lateral preoptic (LPO) nuclei which made them appear over crowded. There are some older reports showing 5-HT projections in hypothalamus. However, some of them had employed older autoradiographic technique and did not cover all the nuclei (Beaudet and Descarries, 1979), while another had used antibody against 5-HT but without the monoamine oxidase inhibitor treatment (Steinbusch, 1981). Although the patterns reported in many of the nuclei were similar to us, we observed higher labelling density in many of the nuclei than their reports because of the superiority of the technique used. None of these previous studies have reported the changes in the labelling pattern in the different hypothalamic nuclei across the rostro-caudal axis.

Some earlier studies have reported the presence of serotonergic cell bodies in the dorsomedial hypothalamic nucleus of rats (Fuxe et al., 1968) (Beaudet and Descarries, 1979) but we did not detect any such cell bodies in hypothalamus. We think that this might be a technical limitation in autoradiographic technique where some non-specific neurons might have absorbed infused 5-HT. A study employing antibody against 5-HT in guinea pig brain also did not detect such cell bodies in hypothalamus (Warembourg and Poulain, 1985).

The ventromedial part of the SCN was one of the highest labelled nuclei in the whole brain. The finding corroborates with results from other rodent models such as hamster (Legutko and Gannon, 2001) and rat (Steinbusch, 1981), and is also consistent with humans (Borgers et al., 2014). According to the earlier tracing studies, SCN receives very dense projections from the median raphe (MR), but not from the DR (Meyer-Bernstein and Morin, 1996) (Muzerelle et al., 2016). Therefore, it can be hypothesized that depletion of 5-HT in the MR or the destruction of 5-HT fibers restricted to the SCN could affect the circadian rhythm. Some previous reports have shown that 5-HT activity on the SCN inhibits the effects of light on the circadian system (Meyer-Bernstein and Morin, 1996) (Bradbury et al., 1997). Moreover, it is reported that the release of 5-HT at the SCN follows the daily rhythm and the behavioral state can strongly influence the serotonergic activity in the circadian clock (Dudley et al., 1998). This strong innervation of 5-HT fibers to the SCN suggests the reciprocal connections of the 5-HT and circadian systems and they may have importance for neurodevelopmental and psychiatric disorders such as ASD and mood disorders, respectively (Ciarleglio et al., 2011) (Takumi et al., 2019).

We noticed an interesting innervation pattern in hypothalamic nuclei regulating the hunger and food intake. The arcuate nucleus was one of the least innervated hypothalamic nuclei. The dorsomedial nucleus exhibited a rostro-caudal innervation gradient. The ventromedial hypothalamic and paraventricular nuclei exhibited both rostro-caudal and intranuclear innervation gradients. The lateral hypothalamic area was too crowded because of the ascending forebrain bundle traversing through it. However, how these wide variations in 5-HT-innervation of these nuclei affect the hunger mechanism is yet to be fully understood. Some studies in mice have shown that there is inverse relationship between brain 5-HT and food intake (Lam et al., 2010).

### 4.3 Amygdala

We observed that most of the amygdalar nuclei received SERT-EGFP fibers in high density except the few nuclei such as lateral amygdala (LA), central amygdala (CeA) and medial amygdala (MeA) were comparatively less innervated. Some of the previous studies have reported serotonergic innervation patterns in the amygdaloid complex using various techniques such as autoradiography (Parent et al., 1981), antibodies against 5-HT (Steinbusch, 1981) or 5-HT transporter (Sur et al., 1996), in situ hybridization (Bonn et al., 2013) and electron microscopy (Muller et al., 2007). However, most of them had either studied only the selected amygdaloid nuclei or have reported lighter density compared to us which might be due to the technical limitations. However, the common finding was that 5-HT fibers distribute densely to the BLA and BMA but less heavily to the LA and CeA nuclei. This was also consistent with the findings reported in a recent review (Asan et al., 2013). In addition, a recent study also showed that BLA is the most labelled nuclei among the different limbic structures which are in the sequence of BLA > NAc > BNST >HIP > CeA > mPFC (Belmer et al., 2017). An anterograde tracing study has reported that amygdalar nuclei receive serotonergic efferents mainly from the DR and very minor projections from the MR (Muzerelle et al., 2016).

We noticed striking species differences in the distribution patterns of SERT-EGFP fibers. On contrary to the rodent pattern, the fiber densities in non-human primate were in the order of central nucleus > basolateral complex > medial nucleus (O’Rourke and Fudge, 2006) (Zeng et al., 2006), while in the human amygdala sequence followed: cortical and anterior amygdaloid nuclei > basolateral and central nuclei (Storvik et al., 2007). Differential expression of the SERT has been linked to species variation in sensitivity to social cues, vigilance to social threats, risk avoidance, responsiveness to changes in reward contexts and mood (Vallender et al., 2009).

Functionally, the BLA is positioned at the pivotal point in the amygdalar circuitry to modify the information received from LA to CeA nuclei (Amano et al., 2011). Coupling this fact with the presence of very dense 5-HT innervation suggests that it could be the primary amygdalar target for 5-HT neuromodulation of fear and anxiety. This is supplemented by findings from a recent pharmacological study which showed that depletion of 5-HT in the BLA reduces anxiety and fear (Johnson et al., 2015). Similarly, action of oxytocin on the MeA nuclei in facilitating the social recognition (Ferguson et al., 2001) could be coupled with the presence of heavy 5-HT innervation in MEAad as the common target for both in modulating the social recognition. Additionally, the decrease in density of 5-HT axons in the CeA and BLA and in the CA3 of the hippocampus due to postnatal social isolation has been linked to depression (Kuramochi and Nakamura, 2009).

### 4.4 Basal Ganglia

We observed fibers in higher density in globus pallidus (GP) compared to caudo-putamen (CP) which contrast with the findings of earlier immunohistochemical studies where no difference in the distribution density was reported (Steinbusch, 1981). Within the CP, fiber densities were higher either ventrolaterally or ventromedially depending on the brain levels which was congruent with a similar study (Mori et al., 1985). Similar to an earlier study (Ternaux et al., 1977) we observed that the innervation density in the striatum increases rostro-caudally. The distribution density of SERT-EGFP fibers demonstrated in our study was consistent with the amount of 5-HT detected in different parts of striatum by liquid chromatography (Beal and Martin, 1985). Similarly, we noticed higher SERT-EGFP fiber density in the external segment (GPe) than the internal segment (GPi) of GP. This pattern was consistent even in primates (Eid et al., 2013). Studies have reported that the innervation pattern in striatum is shared in both rodent and non-human primates (Mori et al., 1985). A recent tracing study showed that striatum and GP receive projection mainly from the supralemiscal (B9) group and minorly from the DR and MR (Muzerelle et al., 2016).

The high serotonergic innervation in the GP suggests that it might be the pivotal site for the 5-HT involvement in the basal ganglia functions or pathophysiology, although it is yet to be confirmed. However, a study has shown that 5-HT influences the reward seeking circuitry involving GPi and lateral habenula (Hong and Hikosaka, 2008). Similarly, the result from one of the studies showing the preferential loss of striatal SERT fibers in Parkinson disease (PD) (Kish et al., 2007) indicates the involvement of 5-HT along with dopamine in the PD pathophysiology. A recent study showed that 5-HT affects the synaptic signaling at thalamostriatal inputs (Cavaccini et al., 2018) which suggests that striatal-dependent functions may be subjected to serotonergic modulation.

### 4.5 Cortex

We found densely concentrated SERT-EGFP fibers in the alternating layers of cortical areas such as mPFC, insular cortex, somatosensory cortex, piriform and rhinal cortices. This was in contrary to the previous immunohistochemical report where they mentioned the uniform distribution of fibers across all the cortical layers (Lidov et al., 1980). Similar to us, one of the immunohistochemical studies reported that the elaborated radial plexus innervates the upper layers and the tangential fibers course through the deeper layers of prelimbic cortex (Miner et al., 2000). We noticed that prelimbic cortex that projects to the limbic areas received higher projection fibers compared to non-limbic regions (i.e. ACC). This could possibly explain the underlying role of 5-HT on the emotional, behavioral and cognitive functions. Any alteration could possibly subserve the pathophysiology of many mental disorders. For instance, reductions in the length of SERT fibers in the orbitofrontal cortex have been found in the postmortem brain of the patient with major depressive disorder (Rajkowska et al., 2017). Similarly, reduced SERT-ir fiber density in the medial frontal cortex, midbrain, and temporal lobe areas have been reported in the autistic brain (Makkonen et al., 2008). A study that employed radiographic techniques reported the prelimbic and rostral agranular insula as the most densely labeled cortical areas sites (Audet et al., 1989). However, in our case we noticed that some other areas like somatosensory (layer 5) and rhinal cortices were equally or even more densely innervated compared to PLA and AI. Anterograde tracing study has shown that both DR and MR projects to prefrontal cortex with DR being the primary source (Muzerelle et al., 2016).

Next, we noticed species difference in the cortical layer labelling density. In our rodent model, a lightly labelled layer 2/3 intervene between moderately labelled superficial and deeper layers of the visual cortex. In the adult cat visual cortex, dense fibers were distributed within the layers I and III (Gu et al., 1990). In monkey, layer IV was the highest labelled layer visual cortical layer, whereas other layers received either less dense or very sparse labelling (Morrison et al., 1982).

Similarly, we observed dense projection within the barrel field area of the somatosensory cortex suggesting its significant role in modulating sensory information. Developmental studies have already demonstrated that 5-HT plays a significant role in the development of barrel formation in cortex (Persico et al., 2000). Not limited to the barrel field, compelling evidence suggests that 5-HT is necessary for the maturation, dendritic arborization, migration, differentiation of many different kinds of neurons and interneurons which are essential for the refined organization of the cerebral cortex (Vitalis et al., 2007).

## 5. Conclusion

In conclusion, we mapped the distribution pattern of the serotonergic neurons across the whole brain region using SERT-EGFP model mice. The use of transgenic animal helped to elucidate the topography of serotonergic system in much greater detail. We identified higher density of projections than previously reported and observed that the densities and pattern changes along the rostro-caudal brain axis. Although serotonergic fibers were ubiquitously distributed in the brain, there was a defined topographic organization of these projections with strikingly high projections in some specific targets. With a couple of exceptions, many of the nuclei with high serotonergic projections were anatomically linked to forebrain limbic structures, suggesting that 5-HT has modulatory effects on emotional and cognitive behaviors. A detailed analysis of the topographical distribution of these neuronal populations will provide an anatomical basis to postulate the physiological role of 5-HT in different behavior and in the understanding of possible alteration in numerous mental and psychiatric disorders.

## Acknowledgement

The authors express sincere thanks to Tsuyoshi Toya, Jun Nomura and Takeshi Kaizuka for their useful help and discussions and all the technical staff of the Takumi Lab for their technical assistance.

## Authors contribution

All authors together conceived and designed the experiments. JRA drafted the manuscript, collected the data and carried out the analysis with advice from KT and TT. All authors were involved in the discussion and editing of the final manuscript. TT supervised the project.

