## Supplemental FIgure 1 and 2 for "Comprehensive Topographical Map of the Serotonergic Fibers in the Mouse Brain"

**Funding information**

This work was supported partly by KAKENHI (16H06316, 16H06463) from Japan Society for the Promotion of Science and Ministry of Education, Culture, Sports, Science, and Technology, Intramural Research Grant for Neurological and Psychiatric Disorders of NCNP, the Takeda Science Foundation and Smoking Research Foundation. Awasthi has been awarded the International Program Associate (IPA) fellowship from the RIKEN - Saitama University joint frontier program, Japan.

For Peer Review

**Abstract**

It is well established that serotonergic fibers distribute throughout the brain. Abnormal densities or patterns of serotonergic fibers have been implicated in neuropsychiatric disorders. Although many classical studies have examined the distribution pattern of serotonergic fibers, most of them were either limited to specific brain areas or had limitations in demonstrating the fine axonal morphology. In this study, we utilize transgenic mice expressing GFP under the SERT promoter to map the topography of serotonergic fibers across the rostro-caudal extent of each brain area. We demonstrate previously unreported regional density and fine-grained anatomy of serotonergic fibers. Our findings include: 1) SERT fibers distribute abundantly in the thalamic nuclei close to the midline and dorsolateral areas, in most of the hypothalamic nuclei with few exceptions such as the median eminence and arcuate nuclei, and within the basal amygdaloid complex and lateral septal nuclei, 2) the source fibers of innervation of the hippocampus traverse through the septal nuclei before reaching its destination, 3) unique, filamentous type of straight terminal fibers within the nucleus accumbens, 4) laminar pattern of innervation in the hippocampus, olfactory bulb and cortex with heterogenicity in innervation density among the layers, 5) cortical labelling density gradually decreases rostro-caudally, 6) fibers traverse and distribute mostly within the gray matter, leaving the white fiber bundles uninnervated, and 7) most of the highly labelled nuclei and cortical areas have predominant anatomical connection to limbic structures. In conclusion, we provide novel, regionally specific insights on the distribution map of serotonergic fibers using transgenic mouse.

**Keywords:** Serotonin, SERT, transgenic mouse, whole brain mapping

For Peer Review

**Supplementary figure 1: Representative images of the innervation density**

**(A)** Innervation density scale 0. olfactory nerve (arrow). (**B**) Innervation density scale 1. ventrobasal thalamic nuclei (circle) **(C)** Innervation density scale 2. Cortex (arrow) **(D)** Innervation density scale 3. Cortex (arrow) **(E)** Innervation density scale 4. CA3 stratum oriens of hippocampus (arrow) **(F)** Innervation density scale 5. Lateral dorsal thalamic nuclei **(G)** innervation density scale 6. Basal amygdaloid complex (arrow).

**Supplementary Figure 2. Schema of the findings in this study**

The main ascending forebrain bundle (blue line) arising from the midbrain raphe cell clusters (B5-B9) branched to give various collaterals (green lines) which ultimately innervate the whole brain, finally to terminate in the olfactory bulb. We analyzed the density and pattern of innervation rostro-caudally throughout the forebrain and cerebellum. Words or abbreviations with asterisks (*) are the most labelled parts within the respective brain region or sub-regions. Specifically, glomerular (gl) layer within the olfactory bulb, globus pallidus (GP) within the basal ganglia, lateral septal nuclei rostral part (LSr) within the septum, suprachiasmatic nuclei (SCN) within the hypothalamus (HYP), basal amygdaloid complex within the amygdala (AMY), dorsolateral and midline

For Peer Review

area within the thalamus (TH), stratum lacunosum moleculare (slm) within the hippocampus (HIP) and deep nuclei within the cerebellum were the most labelled parts. Unique terminal fibers were found distributed within the nucleus accumbens (NAc). The innervation density of cortex (CTX) decreased rostro-caudally. The fibers were arranged in the laminar pattern within the olfactory bulb, hippocampus and cortex.

For Peer Review

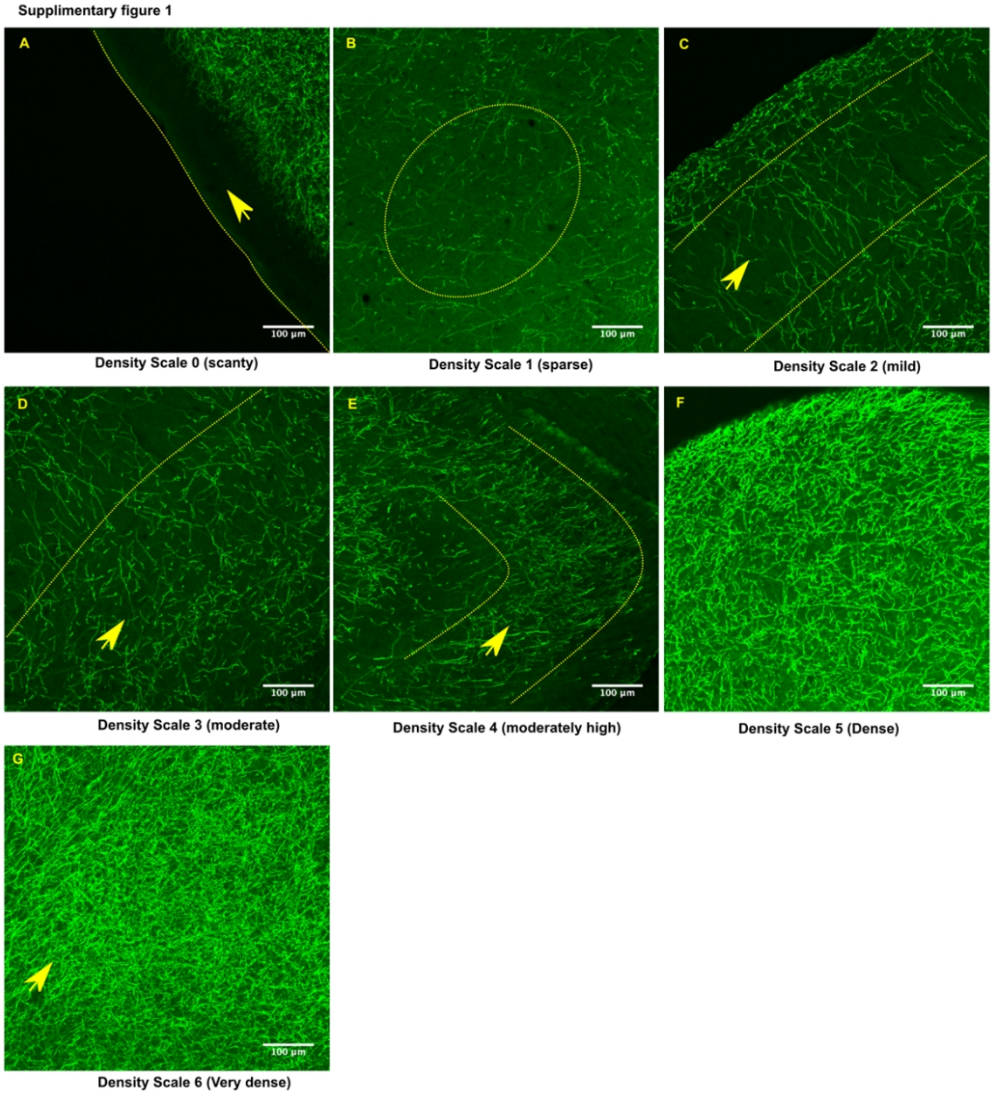


100x111mm (299 x 299 DPI)

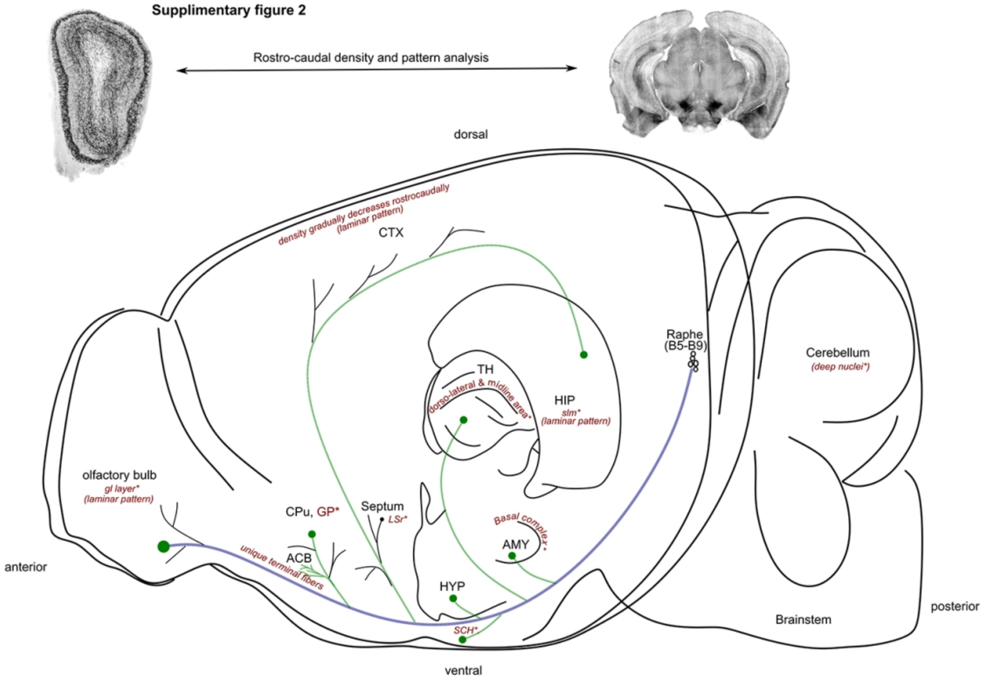


120x84mm (300 x 300 DPI)
